# Long-read powered viral metagenomics in the Oligotrophic Sargasso Sea

**DOI:** 10.1101/2022.09.19.508504

**Authors:** Joanna Warwick-Dugdale, Funing Tian, Michelle Michelsen, Dylan R Cronin, Karen Moore, Audrey Farbos, Lauren Chittick, Ashley Bell, Holger H Buchholz, Rachel J Parsons, Ahmed A Zayed, Michael J Allen, Matthew B Sullivan, Ben Temperton

## Abstract

In the summer months, the waters of the Sargasso Sea are nutrient-limited and strongly stratified, serving as a model system for the predicted warmer and nutrient-limited oceans of the Anthropocene. The dominant microorganisms of surface waters are key drivers of the global carbon cycle. However, the viruses of the Sargasso Sea that shape these host communities and influence host biogeochemical function are not well understood. Here, we apply a hybrid sequencing approach that combines short- and long reads to survey Sargasso Sea phage communities via metagenomics at the viral maximum (80m) and mesopelagic (200m) depths. Taxonomically, we identified 2,301 Sargasso Sea phage populations (~species-level taxonomy) across 186 genera. Over half of the phage populations lacked representation in other global ocean viral metagenomes, whilst 177 phage genera lacked representation in phage isolate databases. Viral fraction and cell-associated viral communities captured in short-read data were distinct and decoupled at both depths, possibly indicating low active lytic viral replication in the Sargasso Sea, with viral turnover occurring across periods longer than the sampling period of three days. Inclusion of long read data was critical for (1) the identification of 79 ecologically important and common viral genomes; (2) capturing the extent of viral genome microdiversity; and (3) enabling the recovery of hypervariable regions in viral genomes predicted to encode proteins involved in host recognition, DNA synthesis and DNA packaging. Host prediction was only possible for ~4% of viral populations. Genomes of phages known to infect *Prochlorococcus* and *Pelagibacter* were poorly represented in our data, supporting recent evidence of low infection levels in the dominant bacterial taxa of oligotrophic regions.

**Subjects:** Bioinformatics, Genomics, Marine Biology, Microbiology, Virology

**Sequence data accession numbers:** PRJNA767318

## INTRODUCTION

Within the North Atlantic sub-tropical gyre bounded by ocean currents, the Sargasso Sea is a vast nutrient-limited (oligotrophic), indigo-blue desert (Menzel & Ryther, 1959). As the site of the world’s longest-running, continuous oceanic study, the physical and bulk biochemical factors that structure this region of open ocean have been well reported (reviewed by (Steinberg et al., 2001): Seasonal changes in surface heat flux and wind stress drive strong thermal stratification of the upper ocean during summer, cutting off nutrient influx from deeper waters and decreasing primary production as nutrients are consumed. Annual vertical mixing of the water column down to ~200m in the winter brings nutrients into the euphotic zone and drives high rates of primary productivity (spring blooms), establishing a robust annual succession of different microbial communities that wax and wane as these nutrients are depleted (Giovannoni & Vergin, 2012). The Sargasso Sea serves as a natural experiment for how increasing temperatures affect microbial community dynamics and associated carbon cycling (Lomas et al., 2013). As oceans warm and become more stratified, oligotrophic systems will expand to encompass more of the ocean surface (Capotondi et al., 2012; Baxter, 2016).

Bacterioplankton communities are the primary drivers of the global carbon cycle (Falkowski, Fenchel & Delong, 2008), and studies of community composition in the Sargasso Sea have provided valuable insights towards understanding productivity and nutrient cycling on the global scale (Venter et al., 2004; Carlson et al., 2009). Sargasso Sea microbial communities are distinct across seasonal and depth gradients (Treusch et al., 2009). Depth-specific niches are occupied by ecologically important heterotrophs, such as Pelagibacter (SAR11), Chloroflexi (SAR202), Deltaproteobacteria (SAR324) and Marinomicrobia (SAR406) (Giovannoni, 1990); (Giovannoni et al., 1996; Gordon & Giovannoni, 1996); (Gordon & Giovannoni, 1996); (Wright et al., 1997) and the phototrophs *Prochlorococcus* ((Moore & Rocap, 1998) and *Synechococcus* (Lomas et al., 2013). *Prochlorococcus* and SAR11 (Pelagibacter) are the dominant phototrophs and heterotrophs in epipelagic surface waters, respectively (Treusch et al., 2009); (Wang & Malanotte-Rizzoli, 2014). The extremely low levels of nutrients in the Sargasso Sea photic zone have constrained the diversity of the microbial communities, as revealed by 16S rRNA community profiling (Treusch et al., 2009). However, metagenomic surveys revealed fine-scale, intraspecific diversity of *Prochlorococcus* and SAR11 (Venter et al., 2004), comprising co-existing populations (Kashtan et al., 2014)(Wilhelm et al., 2007).

Within these populations, distinct ‘core-’ and ‘flexible’ genes/regions, as well as ‘hypervariable regions’ (HVRs), also known as genomic islands(Coleman et al., 2006) between ecotypes are thought to facilitate niche-differentiation within these taxa (Wilhelm et al., 2007; Kashtan et al., 2014; López-Pérez et al., 2020). HVRs are identified when short reads from metagenomic datasets are mapped against genomes from the same environment to reveal regions of very low read recruitment indicating a putative HVR (Rusch et al., 2007).

Bacteriophages are major drivers of both biogeochemical cycles and selection of ecotypes. Ocean viruses directly influence availability of carbon via host lysis (Suttle, 2007); structure microbial communities through negative density dependent selection (Rodriguez-Valera et al., 2009); and alter biochemical function through co-evolution (Lindell et al., 2004, 2007; Scanlan et al., 2015) and metabolic hijacking/reprogramming(Forterre, 2012; Howard-Varona et al., 2020); reviewed in (Breitbart et al., 2018; Warwick-Dugdale et al., 2019a). Viral abundance was identified as a strong predictor of carbon flux from oligotrophic regions in global ocean surveys using ecological network modelling approaches (Guidi et al., 2016). In the Sargasso Sea, abundance of virus-like particles has seasonal and depth-associated structure, with a maximum concentration observed at 80m (Parsons et al., 2012). Correlations between host abundance and total viral particle counts suggested that viral abundance was positively correlated with the abundance of *Prochlorococcus* and negatively correlated with SAR11 abundance (Parsons et al., 2012), potentially indicating that viral communities were dominated by cyanophages, with viruses infecting SAR11 (pelagiphages) poorly represented. Curiously, pelagiphages have been reported as globally ubiquitous and abundant (Kang et al., 2013; Zhao et al., 2013; Martinez-Hernandez et al., 2019; Zhang et al., 2020), but do not contribute significantly to the variance in virally-associated carbon export to depth in the oligotrophic ocean, which is dominated by phages infecting *Synechococcus*) (Guidi et al., 2016). This paradox prompts the question: How are abundant viruses of globally dominant heterotrophs both insignificant in terms in carbon export and contribution to the restructuring of cellular populations in the Sargasso Sea?

One hypothesis of the disconnect between host turnover and viral abundance in SAR11 is that pseudolysogeny or chronic infection is more prevalent than lysis in SAR11 host-virus dynamics, as suggested by low transcriptional activity of pelagiphages in a temperate coastal system (Alonso-Sáez, Morán & Clokie, 2018). Alternatively, due to the enormity of SAR11 populations, a small proportion of susceptible cells within a much larger population of resistant cells could sustain large pelagiphage populations, as long as susceptible hosts possessed some ecological advantage over resistant conspecifics, as observed in *Synechococcus* (Waterbury & Valois, 1993). More recently, observations of very low levels of *Prochlorococcus* infection (0.35–1.6%) in oligotrophic waters replete with cyanophages has been hypothesised to result from a combination of host resistance, low phage adsorption rates and rapid loss of infectivity of virions (Mruwat et al., 2020).

To date, our understanding of the structure of the viral communities in the Sargasso Sea have been limited to seasonal patterns in abundances of viral-like particles (Parsons et al., 2012) and genomic analysis of isolated phages (Sullivan et al., 2005, 2010; Kelly et al., 2013; Zhao et al., 2013; Buchholz et al., 2021a). Here, we used metagenomics to characterise the viral communities at 80m and 200m in both the cellular and viral fractions of the stratified Sargasso Sea during a four-day cruise in July 2017. To maximise recovery of important viral populations, long read metagenomics of the viral fraction was used to overcome issues of assembly fragmentation due to microdiversity and to improve recovery of virally encoded HVRs to facilitate evaluation of their role in niche-adaptation (Martinez-Hernandez et al., 2017; Roux et al., 2017; Warwick-Dugdale et al., 2019b; Zablocki et al., 2021). We compare viral abundance between paired cellular and viral fractions to show that the composition of the viral fraction population did not reflect that of the associated cellular fraction. Viral communities were distinct between depths and comprised many viral populations that were undetected in previous global ocean viral surveys. Phages known to infect SAR11 and *Prochlorococcus*, from both isolates and metagenomic viral contigs where host could be determined, were poorly represented in the viral fractions, suggesting low viral contribution to cellular turnover and nutrient recycling in these taxa.

## MATERIALS & METHODS

### Collection and DNA extraction of host communities and viral assemblages

Metagenomic samples were collected aboard the *RV Atlantic Explorer* at the Bermuda Atlantic Time Series (BATS; http://bats.bios.edu/) station (31°40’N, 64°10’W) via rosette-mounted Niskin bottles during dusk (~19:00 local time) and dawn (~06:00 local time), from depths of 80 m and 200 m, over a period of four consecutive days from the 8^th^-11^th^ July 2017. Host communities (n = 12) were obtained from 5 L of seawater per sample transferred immediately to clean polycarbonate bottles; the cellular fraction was recovered onto 0.22 μm pore Sterivex filters via positive pressure filtration. Each sample was stored in the dark at −20°C in 1mL of SET buffer (0.75 M sucrose; 40 mM EDTA; 50 mM Tris-base). Within a fortnight of collection, DNA from the cellular fraction (which included host DNA, lysogenic viruses and any free viruses attached to cells) was extracted using phenol-chloroform (Fuhrman et al., 1988); (Giovannoni et al., 1990), resuspended in 10 mM Tris-Cl buffer (pH 8.5), and stored at 255 cl:30920°C. Viral assemblages (n = 12) were obtained via sequential filtration of 20 L seawater per sample, followed by iron chloride flocculation (John et al., 2011), with modifications for prevention of DNA degradation and removal of PCR inhibitors (Warwick-Dugdale et al., 2019b). Briefly, peristaltic pumps and 142 mm rigs were used to remove the cellular fraction via sequential filtering through glass fibre (GF/D: pore size 2.7 μm) then polyethersulfone (pore size 0.22 μm) filters, before flocculation and precipitation of viruses via iron chloride. Iron-bound viral particle flocculate was recovered onto 1.0 μm polycarbonate filters (within 4 hours of collection); filters were then stored in the dark at 4°C. Viruses were resuspended (within 5 months of collection) in ascorbate-EDTA buffer (0.1 M EDTA, 0.2 M MgCl_2_, 0.2 M ascorbic acid, pH 6.1), concentrated using Amicon Ultra 100 kDa centrifugal filter units (Millipore UFC910024) and purified with DNase I (to remove un-encapsulated DNA). Viral DNA was extracted using the Wizard^®^ DNA Clean-up System (Promega A7280). In contrast to previous samples derived from a coastal station (Western English Channel site ‘L4’; (Warwick-Dugdale et al., 2019b), Sargasso Sea viral metagenomic DNA required further removal of PCR inhibitors prior to successful sequencing library preparation. This was accomplished using silica membrane spin columns (DNeasy PowerClean Pro Cleanup Kit; Qiagen 12997-50), before elution in 10 mM Tris-Cl buffer (pH 8.5), and storage at 4°C (colder storage temperatures were avoided to prevent freeze-thaw shearing of DNA for long-read sequencing library preparation).

### Library preparation, amplification and sequencing

For short read sequencing, 1 ng of host community DNA and viral assemblage DNA was used to generate Nextera XT libraries (Illumina; manufacturer’s protocol); After amplification (12 cycles) and assessment of library quantity (Qubit; ThermoFisher) and quality (Bioanalyzer; Agilent), library DNA was sequenced as 2 × 300 bp paired-end sequence reads, on a HiSeq 2500 (Illumina Inc.) in rapid mode, by the Exeter Sequencing Service (University of Exeter, UK). In addition, long-read sequences (mean average length ~4 Kbp; Table S1) were generated from viral metagenomic DNA via nanopore sequencing (Oxford Nanopore Technology: ONT) using the ‘VirION’ pipeline (Warwick-Dugdale et al., 2019b), dx.doi.org/10.17504/protocols.io.p8fdrtn). Briefly, ~100 ng of DNA per sample was sheared to ~8 kbp fragments to maximise PCR and sequencing efficiency and amplified using PCR-adapters for Linker Amplified Shotgun Library (LASL) generation. Samples from 200m did not amplify sufficiently for preparation of long-read sequencing libraries, therefore downstream analysis focused on long-read assemblies and analysis of 80m samples. Three long read viral samples from 80m were prepared using the SQK-LSK109 kit and barcoded with native barcoding before being sequenced on a single MinION Mk 1B flowcell (FLO-MIN106; R9.4 SpotON; ONT)

### Sequence processing and assembly

The following bioinformatic pipeline is summarised in Figures S1 and S2. Raw metagenomic short reads from both cellular and viral fractions were initially processed with cutadapt (Martin, 2011) to remove adaptors and PhiX reads (i.e., control library), then error-corrected and quality controlled with bbmap (https://jgi.doe.gov/data-and-tools/bbtools/). Cleaned reads for each sample were assembled independently with metaSPAdes (v3.13.1; (Nurk et al., 2017) using k-mer sizes: 21, 33, 55 and 77. Long-read sequence data was basecalled with high accuracy using Guppy v3.3.0 and demultiplexed with Porechop (v0.4; https://github.com/rrwick/Porechop). Fewer than 42% of the reads were successfully assigned to a barcode, so following removal of adapters and barcodes with Porechop (including removal of reads with adapters in the middle), all reads from the three 80m samples were pooled together and filtered with NanoFilt (De Coster et al., 2018) to remove those with a q-score <10. Pooled, clean, high quality long reads were assembled (via Overlap Layout Consensus: OLC) with metaFlye (Kolmogorov et al., 2020) and Minipolish (including 10 rounds of Racon) (Vaser et al., 2017); (Wick & Holt, 2019), followed by an additional polishing round with Medaka (https://nanoporetech.github.io/medaka/) and two rounds of short-read polishing with cleaned and pooled short reads from matched samples (Stewart et al., 2019).

### Viral sequence recovery and dereplication

To identify Sargasso Sea viruses in our assemblies, contiguous sequences from long and short read assemblies that were circular or ≥ 10kb were processed with VirSorter (v1.0.5; (Roux et al., 2015a) after augmenting its database with the Xfam database of viral HMM profiles from (Guo et al., 2021); contigs resolved into categories 1, 2, 4, and 5 were classed as putatively viral and passed into downstream analyses (Roux et al., 2017). One viral sample (collected July 8th from 200m) was excluded from further investigation due to a lack of viral sequence detection, the result of very low sequencing depth (29.6 Mbp, 100 times smaller than that of other samples). Viral contigs were dereplicated into viral populations by clustering those that shared ≥ 95% nucleotide identity across ≥ 85% of the contig length, using ClusterGenomes (https://github.com/simroux/ClusterGenomes) (Roux et al., 2017). The longest contig within a cluster was selected as the cluster (population) representative. Cluster representatives were used in a second round of clustering with a combined dataset of Global Ocean Viromes 2.0 dataset (GOV 2.0; (Gregory et al., 2019); accession numbers ENA (https://www.ebi.ac.uk/ena): PRJEB402; PRJEB9742; NCBI (https://www.ncbi.nlm.nih.gov/): PRJNA366219) to identify viruses that belonged to known viral populations in Global Ocean datasets and those that were novel to the Sargasso Sea.

### Local and global relative abundance calculations

Next, we investigated which viruses were most abundant in our samples, and how Sargasso Sea viruses are distributed across the Global Ocean. For local abundance calculations, Illumina reads that passed quality controls were competitively recruited to the Sargasso Sea dataset of viral population representatives (derived from both VirION and short-read assemblies) using Bowtie2 (Langmead & Salzberg, 2012), in non-deterministic, sensitive mode; the resulting bam files were parsed in BamM (https://github.comEcogenomics/BamM) to retain reads that mapped at ≥90% read length at ≥95% identity (Roux et al., 2017). The abundance of viral populations within each sample were calculated using mean contig coverage (excluding <5th and >95th percentile – tpmean; (Gregory et al., 2019)using BamM “coverage”. Population representatives with <70% coverage (Gregory et al., 2019) within a sample were assigned an abundance of 0 to minimise false positive detection of populations within a sample (Roux et al., 2017). Coverage values of viral populations were then normalized by total number of reads per metagenome as a proxy for relative abundance.

To investigate the global distribution of the recovered Sargasso Sea viral populations, short reads from this study and from the GOV 2.0 dataset (Gregory et al., 2019) were mapped back to the Sargasso Sea and GOV2 representative viral population contigs. The GOV 2.0 dataset contains 145 samples from five distinct global ecological zones, including the Arctic, Antarctic, bathypelagic, temperate and tropical epipelagic, and mesopelagic. Datasets were subsampled to 5 million reads prior to read mapping to prevent sequencing depth influencing the likelihood of contigs meeting the minimum genome coverage cut-off value (≥70%) and thus possible inflation of the number of rare viruses detected as present in larger datasets compared to smaller datasets. Subsampled reads were recruited against a dereplicated set of Sargasso Sea and GOV2 viral population contigs (using the cut-offs and read recruitment strategy detailed above). Estimated presence/absence values of Sargasso Sea viruses were then calculated singly for each GOV2 site, and for the dataset as a whole.

### Viral classification, microbial survey and host prediction

Having examined the abundance of Sargasso Sea viruses, we next investigated which viruses were present, how the bacterial community was structured, and whether we could predict the hosts of our viral genomes. To classify viruses, open reading frames (ORFs) in viral population representatives were identified with Prodigal (v2.6.1; (Hyatt et al., 2010) in metagenomic mode with default settings. Population representatives were clustered into ICTV-recognized viral genera using vConTACT2 (Bin Jang et al., 2019) alongside RefSeq prokaryotic viral genomes (release 88) for reference to known isolates. Linkage of viral populations to putative hosts was attempted using prophage BLAST, tRNAscan-SE (v1.23) (Chen et al., 2019), and WIsH (v1.0) (Galiez et al., 2017) using a scoring approach previously reported for human gut viromes (Gregory et al., 2020b) (see Supplemental Information).

Recalling previous evidence that at Sargasso Sea, *Prochlorococcus* viruses may be more important to total viral abundance than SAR11 viruses (*sensu* Parsons *et al*., 2012), we evaluated evidence of these phages in Sargasso Sea viromes using two approaches: First, we determined if assembled contigs from Sargasso viromes could be associated with SAR11 or *Prochlorococcus* hosts. The top 100 most abundant viral populations were screened for marker genes associated with cyanophages and pelagiphages using DRAM-v (Shaffer et al., 2020). Specifically, contigs containing photosynthetic *psbA* gene (prevalent in cyanophages) (Sullivan et al., 2006) were extracted as putative cyanophages for manual curation and assessed for completeness with CheckV v0.3.0 (Nayfach et al., 2020). Cultured pelagiphages lack appropriate signature auxiliary metabolic genes which could putatively be used to identify pelagiphages from metagenomic data. However, terminase (*TerL*) genes are commonly used to construct pelagiphage phylogenies (Zhang *et al*., 2020; Buchholz *et al*., 2021) which correlate to clustering of pelagiphage genomes from shared-gene networks (e.g. vConTACT2) (Buchholz *et al*., 2021). To identify pelagiphages, DRAM-v annotations that specified terminase (*TerL*) genes were identified from published pelagiphage genomes, non-pelagiphages (from NCBI refseq, search term: ‘marine terminase in viruses’), *Pelagibacterales* (identified via BLAST hits against terminase from known pelagiphages) and Sargasso Sea viral populations. *TerL* genes were aligned using E-INS-i strategy for 1000 iterations in MAFFT v7.017 (Katoh & Standley, 2016).

Aligned sequences were trimmed with Trimal v1.4.rev15 (Capella-Gutiérrez, Silla-Martínez & Gabaldón, 2009) with sites containing more than 50% gaps removed from the alignment. Alignments were manually checked for overhangs in Geneious v10.2.6 (Kearse et al., 2012). After determining an appropriate substitution model using Model Finder (Kalyaanamoorthy et al., 2017), a phylogenetic tree was constructed using IQ-tree (Nguyen et al., 2015) with rapid bootstrap support generated from 1000 iterations. The phylogeny was visualized in iTOL v5 (Letunic & Bork, 2016), and each clade was subsampled to improve clarity whilst retaining diversity within the tree. Sargasso Sea populations which clustered more closely to non-pelagiphages than pelagiphages were considered to demonstrate dissimilarity to known viruses of SAR11.

Next, we assessed whether genomes similar to those of previously isolated viruses of cyanobacteria and SAR11 were represented in Sargasso Sea viromes through mapping of short reads. High quality short reads were mapped against a dereplicated set of all published cyanophage and pelagiphage isolate genomes (accession numbers: Table S2). Viral population representatives were generated using ClusterGenomes (Roux et al., 2017) as above (cut-offs ≥ 95% nucleotide identity across ≥ 85% genome length). *Escherichia* phage T4 was added as a negative control. Reads were recruited using Bowtie2 (Langmead & Salzberg, 2012), in non-deterministic, sensitive mode, and the resulting bam files were parsed in CoverM (https://github.com/wwood/CoverM) to retain reads that mapped at ≥90% read length at ≥95% identity (Roux et al., 2017). RPKM was calculated as a proxy for relative abundance. To evaluate whether detection of phage isolates was sensitive to minimum genome coverage cutoffs, we repeated this analysis using minimum coverage thresholds of 0-80%.

### Inter-community diversity calculations

Evaluation of the inter-community diversity of the Sargasso Sea viral populations in relation to global viral populations was conducted as follows: Filtered and sequencing-depth normalized read mappings of the complete GOV2 dataset and Illumina reads from BATS against a combined, dereplicated database of Sargasso Sea (this study) and GOV 2.0 (Gregory et al., 2019) viral populations (n=152,979) for abundance calculations (above) were processed to evaluate viral inter-community diversity using the vegan package (Oksanen et al., 2020) (https://CRAN.R-project.org/package=vegan) in R. Viral β-diversity and community structure was evaluated with Principle Coordinates Analysis (PCoA) based on Bray-Curtis dissimilarity (function vegdist;) from cube-root transformed abundance of representative viral contigs. Statistical support for clusters was evaluated using standard deviation and F-tests (function Adonis; 999 permutations), and visualised with ellipses drawn at 95% (inner) and 98% (outer) confidence intervals (function ordiellipse). A Similarity Percentages (SIMPER) analysis (function adonis) was conducted to identify the key Sargasso Sea viral populations driving the community structure depicted via PCoA. To ascertain if the most important contributors to β-diversity were comprised of globally abundant or Sargasso-Sea specific viruses, a bootstrap (n=10000) test was conducted of the median number of Global Ocean Virome (GOV2) samples in which each Sargasso Sea virus appeared (code for bootstrapping available from: ‘https://raw.githubusercontent.com/btemperton/tempertonlab_utils/master/R/StatsUtilities.R’ (Cumming, 2014)).

### Microdiversity calculations

We next evaluated microdiversity in Sargasso Sea virome short- and long-read data to investigate: 1) whether the nutrient-limited waters of the Sargasso Sea were enriched in microdiverse viral populations typically associated with hosts that favour such conditions; 2) whether microdiversity was significantly different between viral populations from 80m and 200m; 3) whether previous findings that long-read viromes capture greater microdiversity in the viral fraction in a coastal region were similarly observed in viromes from nutrient-depleted environments (Martinez-Hernandez et al., 2017; Warwick-Dugdale et al., 2019b). To compare average microdiversity (‘nucleotide diversity’: π; (Nei & Li, 1979) between in Sargasso Sea viruses and those captured in the Global Ocean Virome (GOV2), we used the same approach as Gregory et al. (2019). Long-reads (not available in GOV2 datasets) were excluded from this analysis to avoid potential artefacts associated with sequencing technology. Briefly, all short-read Sargasso Sea viromes were randomly subsampled without replacement to 1M reads using bbmap (https://sourceforge.net/projects/bbmap/). The subsampled reads were assembled, viruses identified, and relative abundances calculated (all methods as above) to generate BAM files. BAM files extracted from Bowtie2 with default parameters were used as inputs to metapop (Gregory et al., 2020a) to call single nucleotide variants (SNVs). Viral populations were only included if ≥ 70% of representative contigs were covered with an average depth of ≥ 10X. SNVs with a quality call of >30 (QUAL score; phred-scaled) were retained, and only those with alternative alleles with a frequency > 1% and supported by ≥ 4 reads were regarded as SNV loci. To minimize sequencing errors and address coverage variations, coverage was randomly subsampled to 10X coverage per locus across the genome. To calculate average microdiversity in Sargasso Sea viruses for comparison to Global Ocean Virome (GOV2) and between depths, mean π from 80m and 200m samples were calculated with 1000 bootstraps of 100 randomly subsampled π values from short-read Sargasso Sea viromes (with replacement). Differences in mean π between depths in Sargasso Sea samples were compared to a null model in which π values from both depths belong to one population (by randomly shuffling of labels and splitting into two datasets of equal size to initial datasets). To investigate whether microdiversity in VirION-derived viral genomes was significantly higher than those in short-read only viral genomes, preliminary work included the generation of additional BAM files by mapping short reads to short-read and VirION assemblies (together; method as above), and the calculation of 100 permuted π values from Sargasso Sea viruses assembled using each approach (i.e., short reads only/VirION pipeline) to generate distributions within those permutations. The permuted percentage increase between the mean π values from each approach was then tested for significance via permutation and bootstrapping test (1000 iterations), as above.

### Investigation of virus hypervariable regions

To investigate the presence and content of hypervariable regions, high quality short reads were mapped to the top 50 most abundant Sargasso Sea viral population representatives (derived from both VirION and short-read assemblies) using Bowtie2 (Langmead & Salzberg, 2012). The resulting bam files were filtered to retain quality mappings (at 95% identity and 70% coverage) using CoverM (https://github.com/wwood/CoverM). The per-nucleotide coverage of the top 50 most abundant viruses was generated from the filtered bam files using bedtools2 (https://github.com/arq5x/bedtools2) and used as input to find HVRs, which were defined as regions that possessed: less than 20% of the whole contig median coverage; at least 600 bp; zones of zero coverage (Mizuno, Ghai & Rodriguez-Valera, 2014). Functions encoded within candidate HVR regions was investigated using a tBLASTx search against the NCBI NR database.

## RESULTS AND DISCUSSION

### Overview

In marine microbial communities, replication is known to be linked to diel cycles, as observed in both photosynthetic- and heterotrophic bacteria, and picoeukaryotes (Jacquet et al., 2001; Ottesen et al., 2014). Assuming that diel cycles would also influence phage production, we sampled cellular and viral fractions for metagenomics over three diel periods to maximise diversity of recovered phage genomes from the Sargasso Sea. DNA was collected from both the “cellular fraction” (>0.2 μm; n = 12; included host DNA, lysogenic viruses and any free viruses attached to cells; 0.22 μm filtration and phenol-chloroform extraction method), and the “viral fraction” (<0.2 μm, n = 12; includes virus-like particles; ferric chloride flocculation and spin column extraction method). Samples were taken at depths of 80 m (the ‘viral particle maximum’, (Parsons et al., 2012)) and 200 m (the mesopelagic), to maximise the diversity of viruses surveyed. DNA recovered from all samples was sequenced to a depth of 12.1 Gbp on the Illumina platform to generate short reads. Additionally, viral-fraction samples from 80 m were sequenced via Nanopore to generate long reads (n = 3, 14.7 Gbp total). Short reads were assembled alone and used for hybrid long- and short-read assembly (‘VirION’; Warwick-Dugdale et al., 2019b) before identification of viral contigs via VirSorter (all bioinformatic processes are described in detail in Methods and Materials and summarised in Supplementary Figures S1 and S2).

### Inclusion of long read data significantly improved recovery of Sargasso Sea viral populations

In total, 3,514 putative viral contigs >10kb in length were recovered from the Sargasso Sea; 2,049 of these were derived from short-read assemblies (12 cellular fraction and 12 viral fraction samples), and a further 1,465 were generated by assembly of VirION reads (three viral fraction samples). Contigs were clustered into 2,301 viral populations (Roux et al., 2017) (equivalent to OTUs). In 1,295 (56%) of the viral populations the longest contig (selected as the cluster representative) was derived from long-read sequencing of pooled 80 m samples. Only 163 (7%) of the viral population clusters contained both long- and short-read derived contigs. No population clusters with two or more members were comprised exclusively of long-read contigs, suggesting either the coverage of the virome in long read data was low, or that short-read sequencing was capturing fragmented genomes across all viral populations for which longer contigs from VirION reads were available. When the 24 short-read assemblies were processed without long reads, 1,044 viral populations were identified, indicating that 55% (1,257) of the total viral populations reported here were only captured with the inclusion of long-read sequencing. Mapping of short-read data to viral population representatives indicated that these viral populations were abundant in the Sargasso Sea (Figure 1A). Representative contigs from long-read assemblies were 38.5% longer than those from short-read assemblies (at median lengths of 17,372 bp and 15,755 bp, respectively; Mann-Whitney U test, *p*<0.001) (Figure 1B). Together, these results suggest that long-read sequencing of viromes enhanced the capture of both different and more complete viral genomes from the Sargasso Sea compared to short-read technology alone.

**Figure 1.**
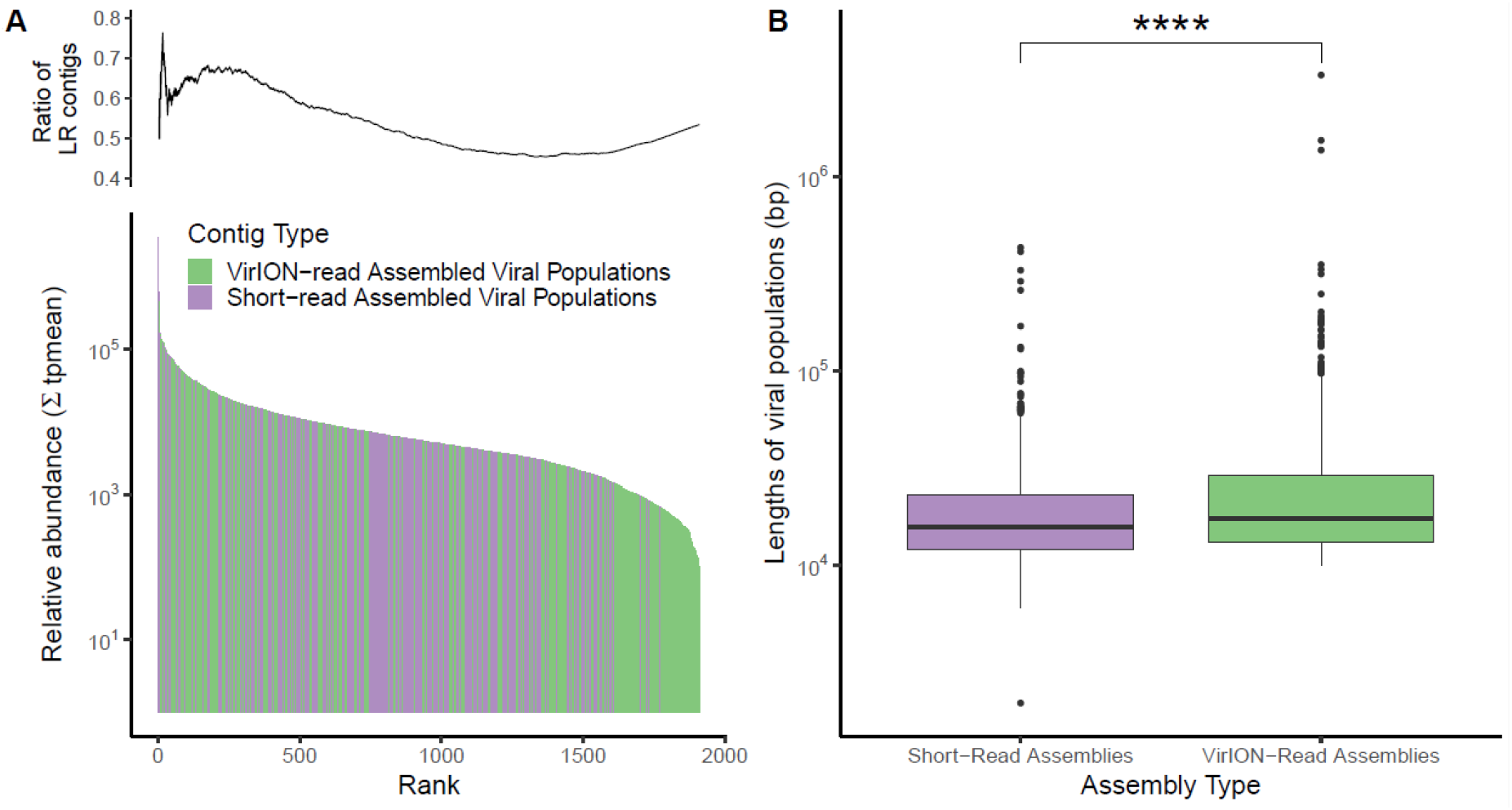
BATS viral populations. **A**. Histogram of sequencing-depth adjusted coverage for viral populations (n = 2,301). Long-read sequencing was able to rescue more viral populations, and these viral populations were abundant. Inset: Cumulative ratio of viral populations derived from long reads. **B** Boxplots showing statistically significant differences in lengths of viral representative contigs assembled from short reads and VirION reads (median lengths 15,755 bp and 17,372 bp, respectively; Mann-Whitney U test, *p* < 0.001).

### Sargasso Sea Viruses form a distinct community

Based on read mapping, 800 (45.7%) of the viral genomes from this study were represented within other subsampled (to 5 million reads) global oceanic viral metagenomes: (GOV 2.0;(Gregory et al., 2019) (Figure 2). Sargasso Sea viral populations from this study appeared more frequently in samples from similarly warm oligotrophic oceanic regions (e.g., TARA_R100000455; Figure 2), and were absent from polar regions, consistent with previously reported ecological patterns of ocean viral communities (Gregory et al., 2019). However, 951 (54.3%) of the Sargasso Sea viral populations were not represented in the global ocean dataset, suggesting that these viruses are either endemic to this subtropical region, or are enriched at this site and below the rate of detection elsewhere. Twenty-four of these ‘endemic/enriched’ viruses were among the top 100 most abundant viruses at this site, indicating their ecological importance in the Sargasso Sea.

**Figure 2.**
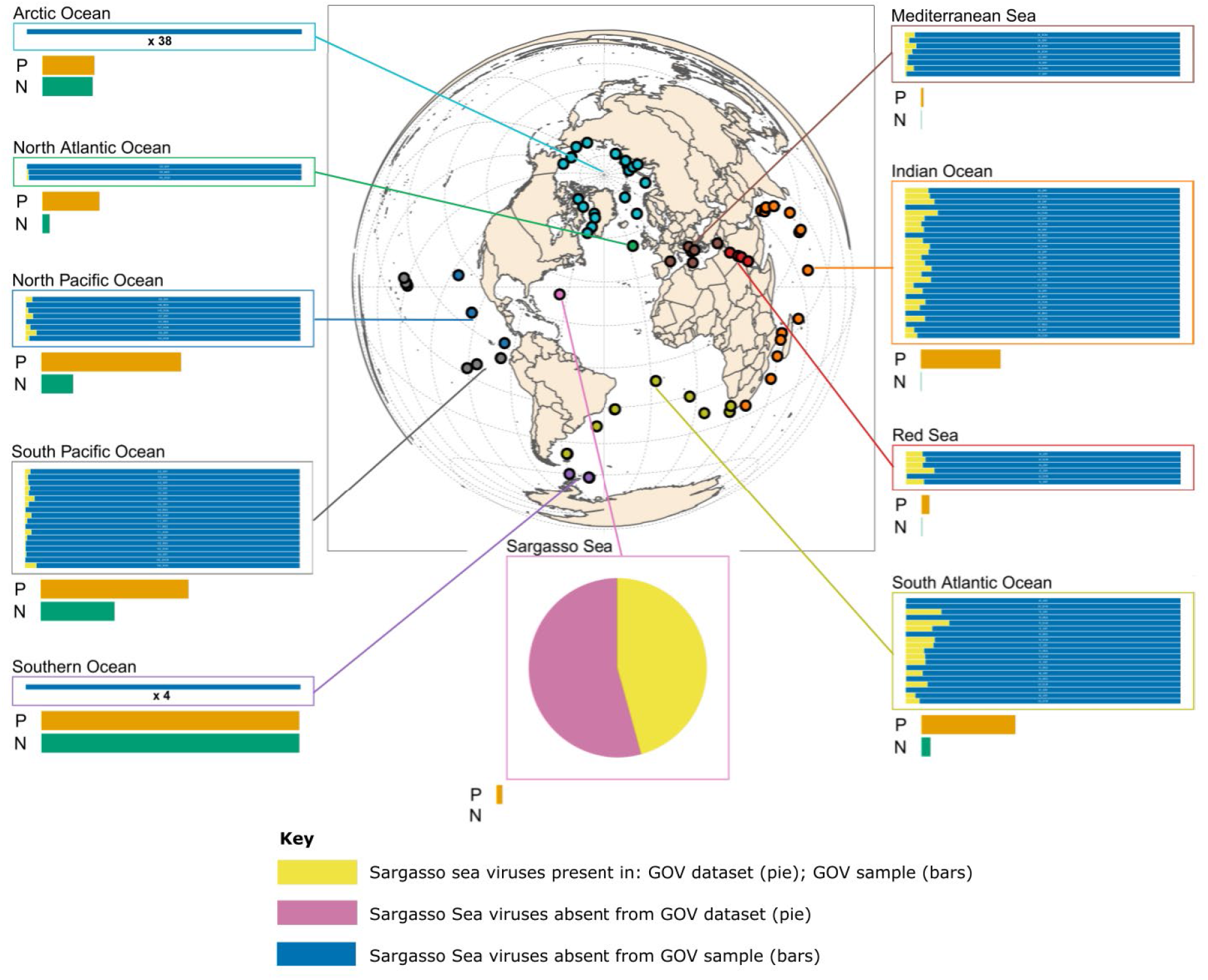
Global distribution of BATS viral populations. Dereplication of the combined GOV 2.0 and BATS dataset resulted in 152,979 viral populations (circular contigs or those with lengths ≥ 10kb). Presence/absence of Sargasso Sea viruses are shown in various oceanic regions (within associated boxes). Under half of Sargasso Sea viruses (45.7%; coloured yellow at all sites) were observed as present elsewhere in the world’s oceans. These viral sequences were identified as present in the GOV 2.0 dataset via competitive read recruitment of subsampled (5 million per sample) reads (mapped at ≥90% read length at ≥95% identity) with >70% genome coverage. The majority of Sargasso Sea viruses were not observed at GOV 2.0 sites (54.3%; coloured pink at the BATS site; coloured blue at other sites), and appear endemic to the Sargasso Sea, (i.e., below the level of detection in the subsampled GOV 2.0 dataset). Levels of nitrogen (NO_3_ = Nitrate+Nitrite^−1^ (umol/kg); ‘N’; coloured green) and Phosphate (PO_4_ = Phosphate^−1^ (umol/kg); ‘P’; coloured orange) were normalised between sample sites (e.g., lowest site median: P=0; highest site median: P=1) and show that more Sargasso Sea virus tend to be present at oligotrophic GOV 2.0 sites.

Inter-community diversity of the Sargasso Sea viral populations was compared to previously established patterns in the Global Ocean Virome (Gregory *et al*., 2019), and revealed that Sargasso Sea viral communities have a distinct community structure (Figure 3). A Similarity Percentages (SIMPER) analysis was performed to determine the key Sargasso Sea viruses contributing to the dissimilarity of this group from viruses from the other ecological zones sampled in the Global Ocean Virome (GOV2) dataset (Figure 4; Figure S2). SIMPER analysis showed that 754 viruses captured in the long-read data explained 9.5% of the variance that discriminated Sargasso Sea viromes from the other viral communities/zones. Viral contigs were classified into two sets (Set A and Set B) depending on their recruitment of viral reads from other GOV2 samples. A set of 80 viral contigs from the Sargasso Sea (‘Set A’) recruited reads from other global viromes and were important in discriminating between temperate and tropical epipelagic (TT-EPI), and temperate and tropical mesopelagic (TT-MES) populations, and between TT-EPI and arctic (ARC) populations. Within these viral populations, 79 out of 80 were comprised of singleton viral populations from long-read assembly that lacked even fragmented short-read assemblies. The median number of Global Ocean Virome (GOV2) samples in which a phage in Set A was observed was 56 (52.5-58 95% CI; out of 142 total). Therefore, we hypothesise that these viruses are common across oceanic regions but missed from existing short-read viral metagenomic datasets. Five of these viral populations had greater global relative abundance than pelagiphage HTVC010P and 51 were more ubiquitous than HTVC010P and other isolates infecting SAR11 and *Prochlorococcus* spp., at middle and lower latitudes (Figure 4). These observations suggest that the VirION approach captured globally distributed and ecologically important viruses that would otherwise have remained unidentified. The remaining 675 viruses (“Set B”), were far less ubiquitous in global oceans, identified in a median of 10 (4.5-17 95% CI) out of 145 GOV2 samples. Four members from ‘Set B’ recruited a large number of reads from at least one site in either the TT-MES or Antarctic biomes (Figure S2), implying some degree of viral import from either upwelling or ocean currents into the 80m and 200m samples. Overall, these results show that the viral community of the Sargasso Sea is distinct in the global ocean and supports the idea that ubiquitous viruses may contribute to the ‘regionalisation’ of viral communities through their relative contribution to overall community structure (Angly et al., 2006).

**Figure 3.**
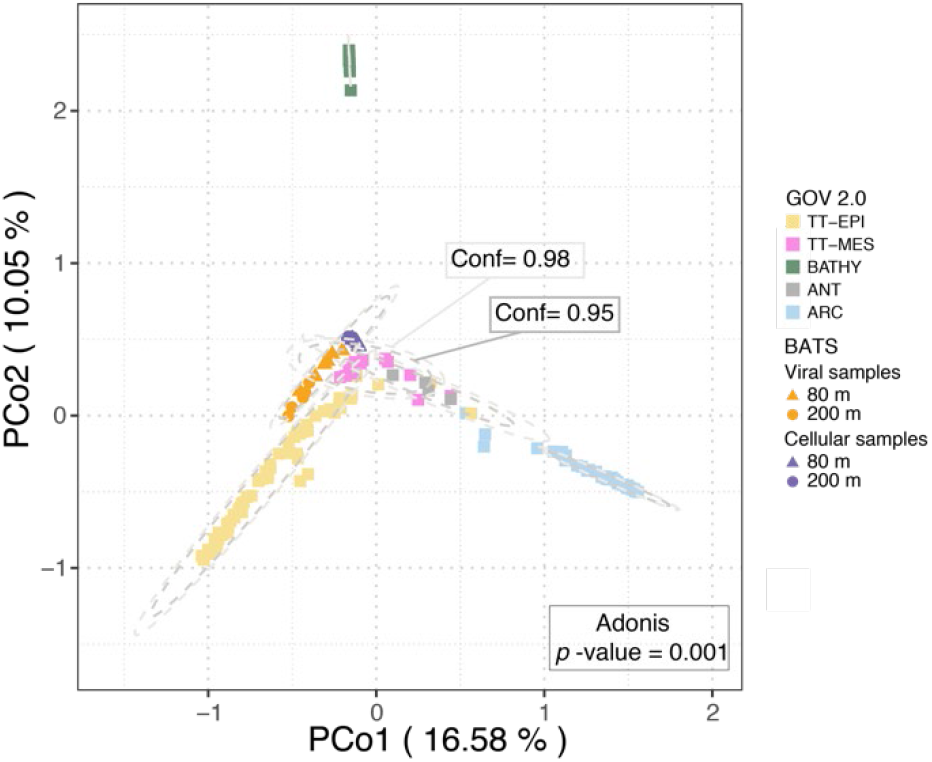
Inter-community diversity of Sargasso Sea and Global Ocean Virome 2 (GOV 2.0) viral populations: Principle coordinates analysis (PcoA) of a Bray-Curtis dissimilarity matrix calculated from mapping GOV 2.0 reads and BATS short reads to a combined dataset of BATS and GOV 2.0 viral populations (contig lengths ≥10kbp); Viral community structure was suggested by ellipses drawn at 95% (inner) and 98% (outer) confidence intervals and analysis of variance (Adonis, *p*-value = 0.001); ARC: Arctic; ANT: Antarctic; BATHY: bathypelagic; TT-EPI: temperate and tropical epipelagic; TT-MES: temperate and tropical mesopelagic.

**Figure 3.**
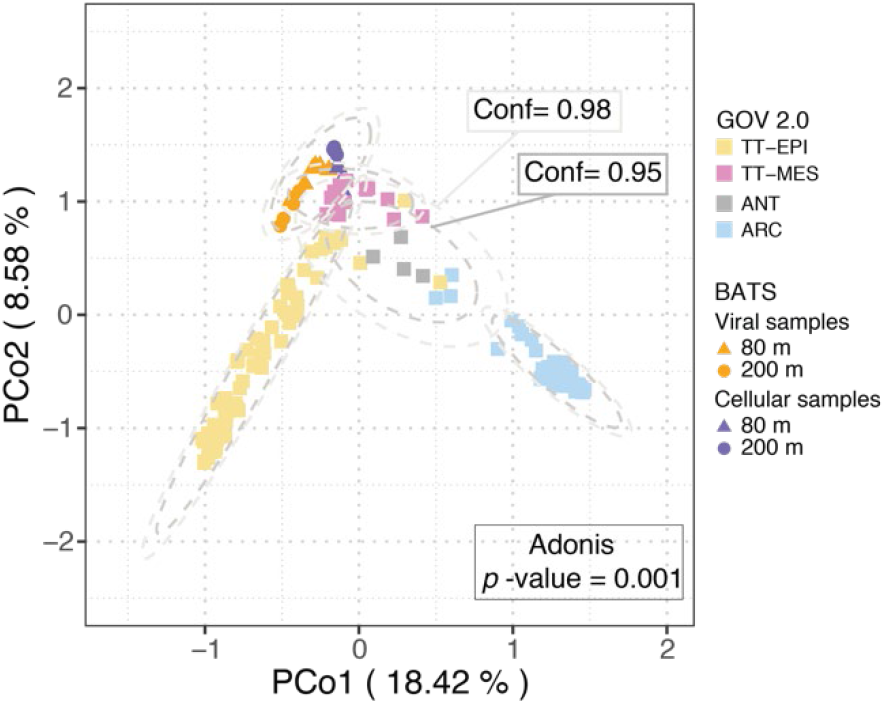
After the removal of BATHY samples.

**Figure 4.**
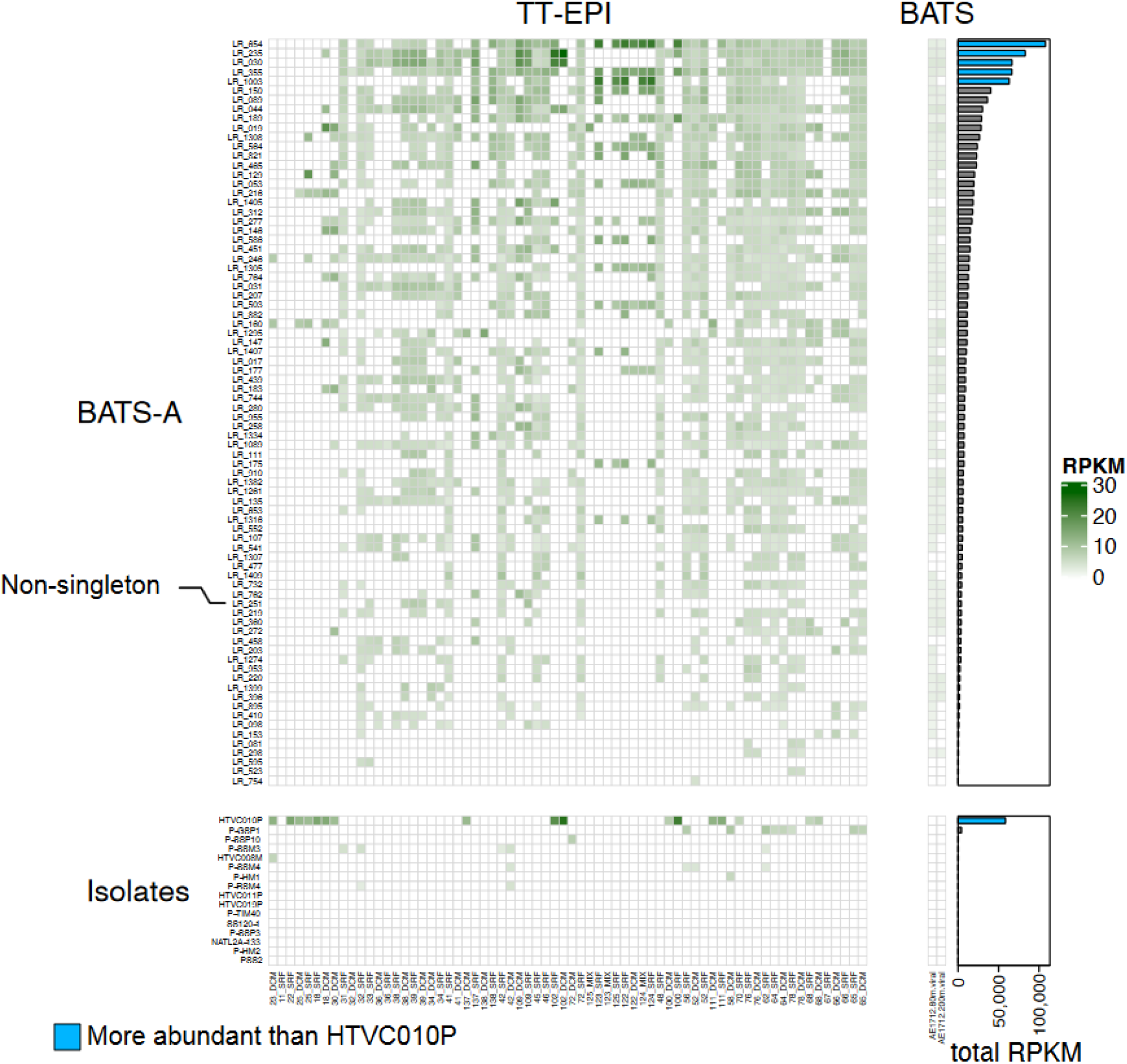
Global relative abundance of Sargasso Sea long-read viral contigs from this study (‘BATSA’) and isolates that infect SAR11 and *Prochlorococcus* (‘Isolates’). BATS-A viruses were important for discrimination between temperate and tropical epipelagic (TT-EPI) and temperate-tropical mesopelagic (TT-MES) viral populations in SIMPER analysis. 79 out of 80 BATSA viruses were recovered from ‘singleton’ populations (i.e., they were not captured by short-read assemblies); five of these viral populations had greater global relative abundance than Pelagiphage HTVC010P in the epipelagic (at middle and lower latitudes). The bootstrapped median (n=10,000) number of Global Ocean Virome (GOV2) samples in which a BATS-A viruses were observed was 56 (52.5-58 95% CI). Thus these viruses are common across oceanic regions, but were missed from existing short-read viral metagenomic datasets.

Sargasso Sea viral populations from both 80m and 200m were most similar to viromes from other temperate and tropical epipelagic (TT-EPI) regions (Figure 3). Three GOV 2.0 viromes (station 155_SUR, station 72_MES, station102_MES) were removed since they were outliers in both PCoA and Shannon’ H analyses (Gregory et al., 2019; Figure S3). The similarity of the Sargasso Sea 200m samples to other TT-EPI (sampling depth: 0-150m) samples, rather than other temperate-tropical mesopelagic (TT-MES) samples (sampling depth: 150-1000m) is concordant with the nature of the Sargasso Sea as an oligotrophic subtropical gyre system where sunlight can penetrate to greater depths (e.g., 1% of surface blue light at 149 m; Clarke, 1936) than in nutrient rich, productive waters. This observation is further illustrated by the increased depth of the Deep Chlorophyll Maximum at BATS during sampling (100m) compared to other regions sampled during the Tara Oceans expedition (South Atlantic Ocean: 40m; Indian Ocean: 25-80m; Mediterranean Sea: 50-70m; Red Sea: 60-80m; Brum et al., 2015). In fact, the Sargasso Sea at 200m may not be regarded as intrinsically mesopelagic; a better name might be the altum- (deep)-epipelagic. Thus, depth seems at best a rough proxy for changes in environmental parameters down the water column, and during summer stratification in the Sargasso Sea depths of >150m may not have the presumed characteristics of oceanic ‘mesopelagic’ regions. Further investigation comparing these results to viral communities sampled after the spring bloom would ascertain whether the same pattern is repeated throughout the year, when vertical stratification is either absent or minimised.

### Viral fraction and cell-associated viral communities are decoupled at both depths

The physical stratification of the ocean due to seasonal warming (and therefore reduced density) of the upper layers, combined with the attenuation of light with depth, has long been understood as an important factor in structuring pelagic, microbial communities down the water column (Morris et al., 2005; Carlson et al., 2009; Treusch et al., 2009; Vergin et al., 2013). Likewise, the viral community composition in the Sargasso Sea differed significantly along the two ecological zones, as delineated by depth (Figure 5a). These results support previous evidence that phages are vertically distributed (Gregory et al., 2019) in the same way as their bacterial counterparts, corresponding to the stratification of the water column during summer (Giovannoni & Vergin, 2012).

**Figure 5.**
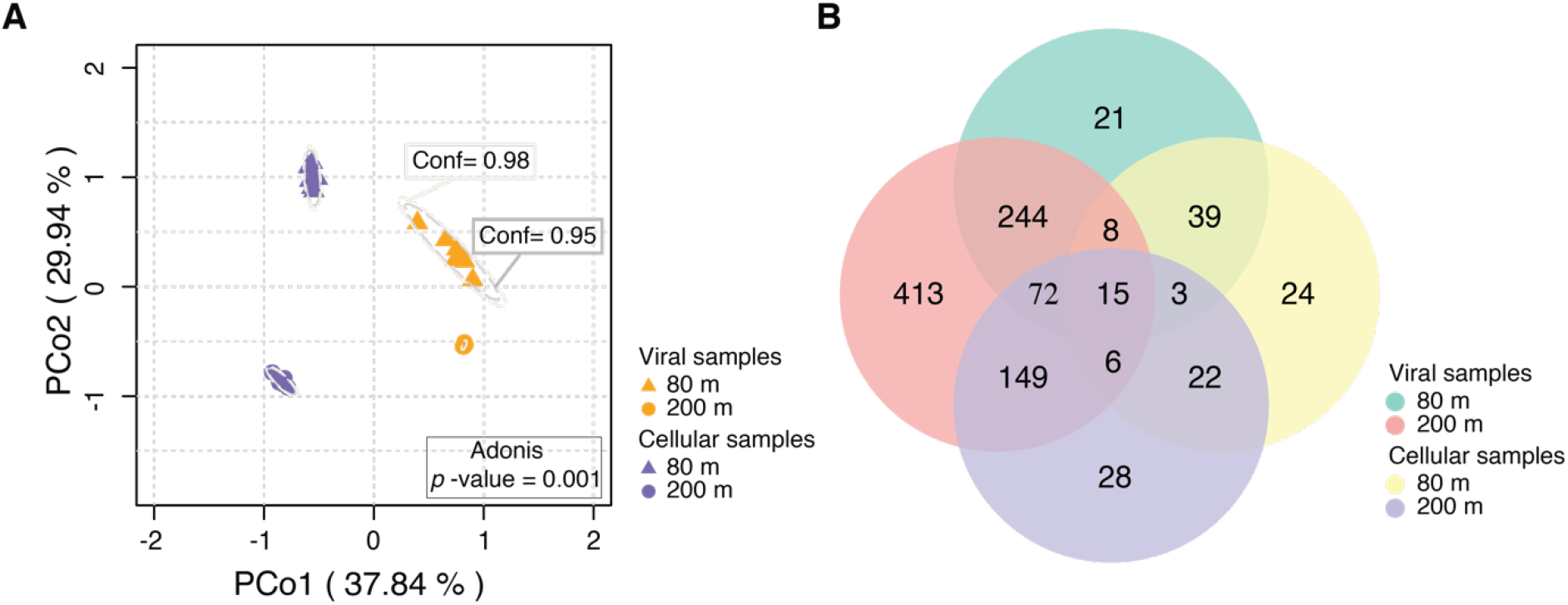
BATS viral populations clustered on the basis of sample fraction type (c = cellular; v = free particulate) and depth (80m; 200m)**; A**. Principal coordinates analysis (PCoA) of a Bray-Curtis dissimilarity matrix calculated from reads mapped to BATS viral populations (n = 2,301) derived from short-read and VirION assemblies. Viral community structure was suggested by ellipses drawn at 95% (inner) and 98% (outer) confidence intervals and analysis of variance (Adnois, *p-* value = 0.001). **B**. Number of viral populations (n = 1044) found in 80m viral samples, 80m cellular samples, 200m viral samples, and 200m cellular samples, as identified from short-read only assemblies.

Curiously, there were significant differences in composition and membership of viruses between cellular and viral fractions from the same depth. Bacteriophage populations in the Sargasso Sea formed discrete ‘cellular fraction’ and ‘viral fraction’ populations, at both the ‘viral maximum’ depth (i.e. 80 m; Parsons *et al*., 2012) and in the mesopelagic (Figure 5a). This zonal partitioning was also observed when long-read derived contigs were excluded from the analysis (Figure S4), indicating that it is not an artefact of sequencing technology. Sixty-five percent of viral populations from the short reads (678) were only detected in the ‘viral fraction’, whereas 7 % (74) were discovered solely in the cell-associated fraction, and 28 % (292) were detected in both fractions (Figure 5b). Previous investigations of soil microbial communities and their phages have examined coupled ‘cellular’ and ‘viral’ fractions of samples for viral genomes and found that the majority (77%) were present in the cellular fraction, despite the samples for each fraction being collected years apart (Emerson et al., 2018; Trubl et al., 2018). However, when viromes and cellular fractions of the same samples were compared, as undertaken here, just 9% of viral populations were shared between fractions, with >90% present in only the viral fraction, and <1% unique to the cellular fraction (Santos-Medellin et al., 2021). Because the Sargasso Sea cellular fraction here may include any free viruses attached to- or caught on cells during the filtration step of our protocol, it is not possible to discount the presence of DNA from free viruses here. The large surface area of filters used to separate cellular and viral fractions, and the relatively low concentration of bacterioplankton cells during the sampling campaign (80m: ~5-7.7 × 10^8^ L^−1^; 200m: ~1.2-3.2 × 10^8^ L^−1^) make it unlikely that viruses were removed from the ‘viral’ fraction by clogging on filters. Thus, the effect of filtration is unlikely to cause the degree of dissimilarity between the free-particle and cell-associated fractions observed in our data. Presence-absence analyses showed that this decoupling of fractions was less pronounced in the 100 most abundant Sargasso Sea viruses (Figure S5), thus rare viruses may be partially driving the dissimilarity in viral- and cellular fraction viral populations. However, Bray-Curtis analysis is robust to the effects of sample size (Schroeder & Jenkins, 2018), so the partitioning effect illustrated via Principal Coordinates Analysis is not likely due to under-sampling of rare viruses.

One explanation for the decoupling of cell-associated and viral fraction populations is that the viral fraction population represents the integral of infections in the cellular fraction over time, whereas the cell-associated populations are a mix of prophages, remnant phages, active lytic infections and a small number of phage particles trapped on filters. Viral turnover in oligotrophic waters has been estimated at 2.2 days, compared to 0.82-1.3 days in coastal waters (Noble & Fuhrman, 2000). Here, sampling was conducted over three days to capture viruses across full host growth and lytic cycles to avoid biases inherent to ‘snap-shot’ surveys, where asynchrony in infection cycles can cause apparent dissimilarities between free and cell-associated viral populations. Decoupling between free and cell-associated viral populations suggests that viral turnover in the Sargasso Sea occurs over periods longer than our sampling campaign. Loss of infectivity and decay of viral particles by sunlight (Suttle & Chen, 1992) could also contribute to decoupling. However, at the depths sampled here (80 m; 200 m), sunlight-induced viral decay is likely to be lessened compared to surface samples (5m) where previous turnover estimates have been calculated (Noble & Fuhrman, 2000)

An alternative explanation for decoupling may be that the rate of active lytic viral replication in dominant members of the community such as SAR11 is low in the Sargasso Sea, as was previously observed in a coastal system (Alonso-Sáez, Morán & Clokie, 2018). This parallels a recent report that high abundances of free cyanophages coincided with low levels of *Prochlorococcus* infection in oligotrophic waters, proposed to result from a combination of loss of infectivity and low adsorption efficiency, alongside host resistance (Mruwat et al., 2020). Success rates of SAR11 cultivation from dilution-to-extinction culturing experiments have been speculated to indicate that a large proportion of streamlined heterotrophs such as SAR11 and OM43 are possibly dormant, and therefore not contributing to viral turnover (Henson et al., 2020). In this scenario, viruses are released into the viral fraction but are unable to efficiently re-infect the cell-associated fraction, decoupling the two populations. High rates of lysogeny in taxa such as SAR11 would produce a similar outcome, although the most abundant pelagiphages in the oceans do not encode known mechanisms of chromosomal integration (Zhao et al., 2013; Martinez-Hernandez et al., 2019). There was no evidence in environmental data collected on the cruise (C. Carlson, pers. comm.) to suggest entrainment of water from outside the gyre, nor do the high numbers of site-specific viral populations in the Sargasso Sea (Figure 2) support possible decoupling by significant entrainment of viruses from outside the sample site. Stable host-virus co-existence and low infection rates in oligotrophic waters imply that phages here are not likely the main control on host abundance, but may instead play an important role in the evolution of clonal, fine-scale diversity in the host population (Waterbury & Valois, 1993). Phages have been shown to increase host diversity on this micro sale in host-phage model systems (Brockhurst, Buckling & Rainey, 2005), and the high divergence in potential phage recognition sites observed in the HVRs of *Pelagibacter* isolates (Rodriguez-Valera et al., 2009) supports the idea that phages drive host microdiversity in the Sargasso Sea.

### Bioinformatic prediction of hosts remains challenging despite inclusion of MAGs

To determine viral taxonomic classification, we clustered the 2,301 Sargasso Sea viral population representatives for genera-level classification based on shared-gene networks using VConTACT2. RefSeq prokaryotic viral genomes (NCBI RefSeq v88 release) were included to assign family-level taxonomy to clusters. Out of 548 viral clusters, 186 contained Sargasso Sea viral sequences, of which 177 lacked a representative sequence in the RefSeq database, consistent with previous studies of environmental viral communities (Angly et al., 2006); (Roux et al., 2015b); (Krishnamurthy & Wang, 2017) (Hurwitz & Sullivan, 2013; (Paez-Espino et al., 2016).

All phages assigned a taxonomy clustered with viruses within the order *Caudovirales* except one genome from family *Microviridae* with the order level *Petitvirales*. Few viral family-level taxonomic annotations were predicted: 26 *Podoviridae;* 3 *Myoviridae*; 3 *Siphoviridae*; 1 *Microviridae*. Among the top 50 most abundant viral populations, 8% were classified, all of which were resolved as either *Podoviridae* or *Myoviridae*, highlighting the need for improved representation of abundant environmental viruses within reference databases (Paez-Espino et al., 2016). We next tried to assign putative hosts to Sargasso Sea viral populations by recovering metagenome-assembled genomes (MAGs) from cellular metagenomes. Assembled contigs were binned using sequence composition, relative abundance, and taxonomical classifications to group contigs into MAGs (Gregory *et al*., 2020). In total, we obtained 89 MAGs with ≥ 70% completion and ≤10% contamination that were used for host-prediction (Table S3). Considering the intrinsic challenges of *in silico* host prediction (Khot, Strous & Hawley, 2020; Moon & Cho, 2021) a scoring matrix was developed to combine the results from prophage blast, tRNA scan, and WIsH to improve the accuracy of host assignments. Despite this, only six out of 2,301 viral populations (0.26%) were successfully linked to three hosts which belonged to the phylum of Actinobacteriota (n = 2) and Chloroflexota (n = 1). Therefore, assigning putative hosts to viral contigs from metagenomes remains a major barrier for understanding environmental viral ecology.

### Evidence indicates low abundance of both cyanophages and known pelagiphages in the Sargasso Sea

During the summer months, cyanobacteria (*Prochlorococcus*) are the dominant phototrophs in the Sargasso Sea, with estimated relative abundances of up to 35% in the euphotic zone (Treusch et al., 2009). Also highly abundant, SAR11 comprises 20-40 % of the total bacterioplankton community in open ocean systems (Morris et al., 2002); (Rappé & Giovannoni, 2003). Previously, total viral abundance was shown to be negatively correlated with SAR11 abundance and positively correlated with *Prochlorococcus* over seasonal scales, leading to the hypothesis that viral communities were dominated by cyanophages with pelagiphages poorly represented (Parsons et al., 2012). Composition analysis of the viral fraction here provides supports a dearth of pelagiphages. Recruitment of reads to *Pelagibacter* phage HTVC010P (a virus previously cited as the most abundant on Earth and isolated from the Sargasso Sea; Zhao et al., 2013), failed to meet the minimum genome coverage to be classified as present, even at a relaxed cut-off of >15% (Figure S7). Moreover, *TerL* genes identified in the 100 most abundant viral populations did not cluster with known pelagiphage terminase genes within a phylogenetic tree (Figure S6). Thus, we found no evidence in either assemblies or short-read data of abundant viruses in the Sargasso Sea that closely related to previously isolated pelagiphages in our samples. Surprisingly, this was also true for cyanophages, in contrast to predictions by Parsons et al (2012). Among the top 100 most abundant viral populations, only four contigs were identified as potential cyanophages via presence of the *psbA* gene, with just two of these having completeness over 80%; these two populations were the 26^th^ and 45^th^ most abundant in the Sargasso Sea. In addition, when short read sequences from both the viral fraction and cellular fractions were recruited directly to a database of all published cyanophage and pelagiphage genomes, only a single isolate genome (*Prochlorococcus* phage P-SSM2, GU071092.1) was identified as present until the minimum genome coverage exceeded >65% (Figure S7).

Rates of successful infection of *Prochlorococcus* in the oligotrophic surface waters of the North Pacific Subtropical Gyre (NPSG) were previously found to be low (Mruwat et al., 2020), despite a high abundance of cyanophage virions (~2.2% of the viral fraction) (Mruwat et al., 2020). The lack of representation of abundant pelagiphages and cyanophages observed in this study suggests similarly low levels of active infection in the Sargasso Sea. Metabolic processes associated with pelagiphage infection were poorly represented in transcriptomic data from a coastal system collected over a 2-year period (n = 8), suggesting that chronic infection was more prevalent than lytic infection (Alonso-Sáez, Morán & Clokie, 2018). This, coupled with predicted dormancy rates of up to 85% in SAR11 cultured isolates (Henson et al., 2020) may further limit efficiency of pelagiphage infection in the Sargasso Sea. In contrast to the NPSG study, we found little evidence of abundant pelagiphages and cyanophages in the viral fraction from the Sargasso Sea. An important difference between these regions is that phosphate concentrations in the Sargasso Sea are up to two orders of magnitude lower than those found in the NPSG (Wu et al.).

Synthesis of viral particles imposes a high phosphate requirement on infected cells and phosphate limitation restricts lytic infection in both cultures of cyanobacteria and natural marine communities (reviewed in (Wilson & Mann, 1991)). Therefore, we propose that in the NPSG, infection of a small proportion of the cyanobacterial community is sufficient to maintain abundant viral particles. In contrast, in the Sargasso Sea, increased P-limitation reduces viral production to a level where known pelagiphages and cyanophages (either isolates or closely related phage) are diluted to below levels of detection with metagenomic approaches. Here, clustering of the *TerL* genes identified in the 100 most abundant viral populations (FigureS3) implies that the most abundant Sargasso Sea viruses captured are most like those that infect copiotrophic hosts (e.g. *Pseudoalteromonas* and *Flavobacterium* phages). This could suggest that boom-bust, particle associated interactions may be of greater importance than previously imagined in nutrient-limited water. However, as a result of the challenges of host-prediction, highlighted by our results here, this hypothesis is highly speculative at present, and more robust methods of host prediction are required for further investigation.

### Sargasso Sea viral populations are microdiverse

Given that the Sargasso Sea bacterial community has more fine-scale, intraspecific diversity than variation at the ‘species’ (or ‘macrodiverse’) level (Venter et al., 2004; Treusch et al., 2009), we determined whether the same could be true of their phages, and how levels of microdiversity might compare to those of global oceanic datasets. Here we report high microdiversity (i.e. intra-population diversity; π (Nei & Li, 1979)) values for Sargasso Sea viral populations across both depths (mean π: 3.411×10^−4^ (2.473×10^−4^ − 4.334×10^−4^, 95% CI) (Figure S8), comparable to those recorded at similar latitudes in the Global Ocean Virome (GOV 2.0; Gregory et al., 2019). The microdiversity of Sargasso Sea viral populations from 80m and 200m samples was not observed as significantly different (permutation and bootstrapping test: *p*=0.164; Figure S9b). This result does not align with those of (Gregory et al., 2019), who reported higher levels of microdiversity in mesopelagic temperate and tropical viromes than those from the epipelagic in the GOV 2.0 dataset. However, it is possible that the altum-(deep)-epipelagic nature of viral communities in the Sargasso Sea at 80-200m works to reduce the signal of increasing microdiversity at depth observed in tropical and temperate viromes (Gregory et al., 2019).

Additionally, we compared microdiversity of Sargasso Sea viral populations obtained using only short reads, and those captured with the inclusion of VirION reads, as previously we have observed that long reads assemble microdiverse viral genomes of Western English Channel viral assemblies better than short-reads due to the benefits of long-read assembly (Warwick-Dugdale et al., 2019b). The average microdiversity values for viruses derived from the two assembly types were 4.213×10^−4^ (3.283×10^−4^ – 5.326×10^−4^, 95% CI) and 2.03×10^−3^ (1.69×10^−3^ – 2.36×10^−3^, 95%CI)., for short-reads and VirION reads, respectively (Figure 6b), which represents an average increase of 389% (264.559 – 551.95%, 95% CI) in the π value calculated for viral genomes captured by VirION compared to those assembled from short-reads (permutation and bootstrapping significance test: *p*<0.001) (Figure S10). Because π values are calculated from short reads which map to viral contigs, rather than the viral contigs themselves, this finding is not influenced by residual error in long-read derived contigs. This result confirms that VirION sequencing facilitates the capture of more viral microdiversity than is possible from short-read sequencing alone. However, the question as to why oligotrophic regions such as the Sargasso Sea produce highly microdiverse viral assemblages remains open. The low infection rates observed in the NPSG (Mruwat et al., 2020) suggests that phages here are not likely the main control on host abundance. Instead, such viruses may play an important role in the evolution of clonal, fine-scale diversity the host population (as proposed decades ago by Waterbury & Valois, 1993). Phages have been shown to increase host diversity on this micro sale in host-phage model systems (Brockhurst, Buckling & Rainey, 2005), and the high divergence in potential phage recognition sites observed in the HVRs of Pelagibacter isolates (Rodriguez-Valera et al., 2009) supports the idea that phages drive host microdiversity in the Sargasso Sea.

**Figure 6.**
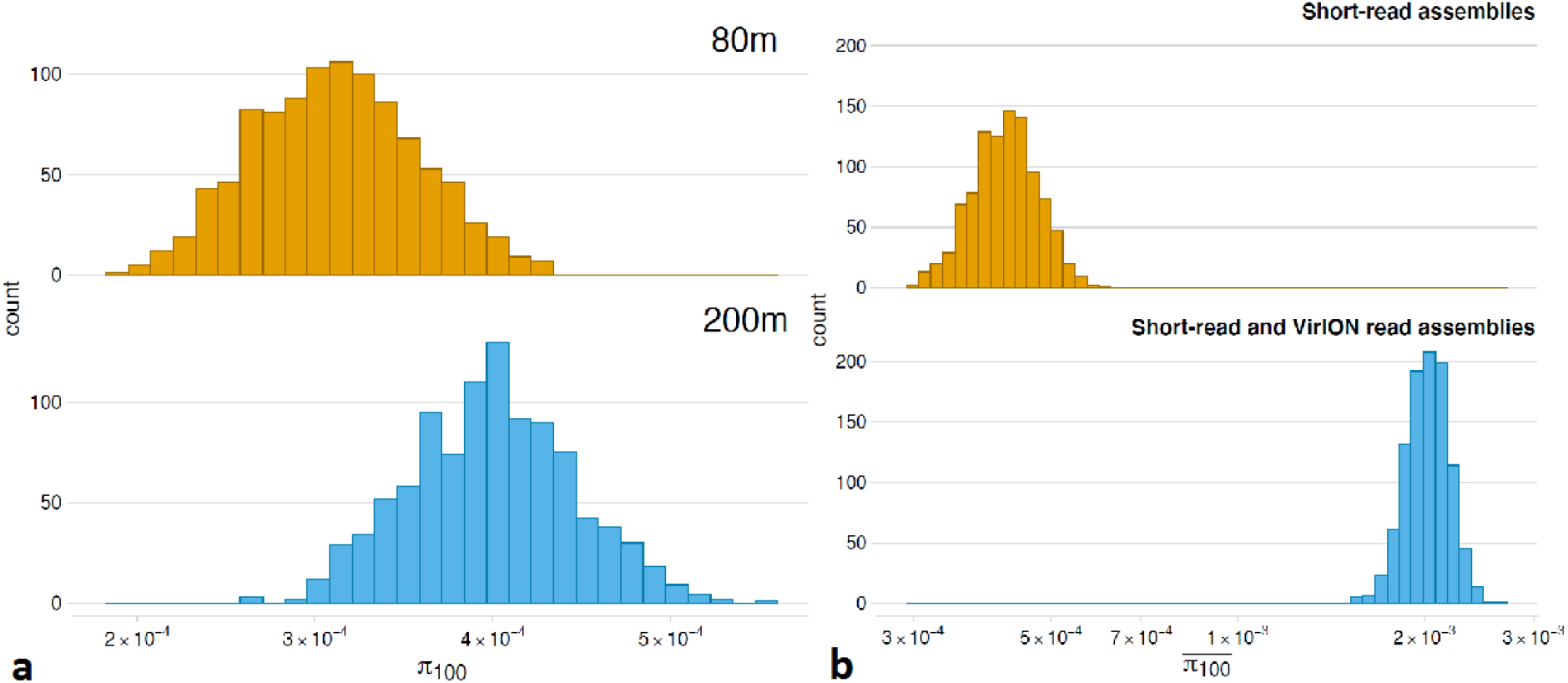
Microdiversity (intra-population diversity; mean π, calculated as (Gregory et al., 2019) of Sargasso Sea viral populations. **A**: The microdiversity of Sargasso Sea viral populations from 200m and 80m samples (mean π: 3.119×10^−4^ (2.347×10^−4^ – 4.094×10^−4^, 95% CI) and 3.967×10^−4^ (3.072×10^−4^ – 4.909×10^−4^, 95% CI), respectively). On average, microdiversity was ~30% greater in viral genomes from 200m than those from 80m samples, but this was not a significant difference (permuted significance test: *p*=0.164; Figure S9b). **b**. VirION reads facilitated the capture of viral genomes with high microdiversity: mean π was 388.668% (264.559 – 551.95%, 95% CI) greater for viral genomes captured by VirION compared to those assembled from short-reads (permuted significance: p<0.001).

Because increased host microdiversity will expand phage niches, phage adaptation to emerging host ecotypes may increase phage microdiversity, as predicted by the ‘Diversity Begets Diversity’ (DBD) theory of species interactions (Calcagno et al., 2017; Madi et al., 2020). DBD, which posits that existing diversity will promote the evolution of further diversity by niche construction (plus other types of interaction), appears to be most prevalent in low-diversity biomes (Madi et al., 2020). Thus, Sargasso Sea host communities with high microdiversity may be predated upon by viruses with correspondingly high microdiversity, providing positive feedback that may magnify the importance of microdiversity in the function of this system. At species level (macrodiversity), bacterial diversity is negatively correlated to the magnitude of Net Heat Flux (NHF; i.e. air-sea flux of heat into an oceanic system) (Smyth et al., 2014). In accordance with the Royal Family Model (Breitbart et al., 2018), the strong positive NHF (Cornillon & Stramma, 1985; Lomas et al., 2011) and associated low macrodiversity in the host community (Treusch et al., 2009) of the summer-stratified Sargasso Sea could constrain viral evolution to fine-scale, nucleotide level changes that maximise niche-filling, whilst increasing microdiversity, amongst the well-established and well-adapted host-communities.

This putative mechanism may explain the strong positive correlation previously observed between viral microdiversity and PAR (Photosynthetic Active Radiation) (Gregory et al., 2019). This idea is corollary to the proposition that increased fine-scale viral microdiversity may decrease more coarse-level diversity by promoting competitive exclusion (Hart, Schreiber & Levine, 2016), proposed to explain regional scale diversity patterns observed at a in the GOV 2.0 dataset (Gregory et al., 2019). Recalling the role of the Sargasso Sea as a model for regions of the ocean that are warming due to anthropogenic climate change, it is clear that this finding has implications for global viral diversity. Viral communities in areas of the ocean that are warming may exhibit decreasing macrodiversity. Further investigation towards a better understanding of what drives viral microdiversity, and how this translates into multiscale host and viral ecology is required.

### Hypervariable regions of Sargasso Sea Viral genomes encode proteins putatively associated with host recognition, DNA synthesis and packaging

Like their prokaryotic hosts, phage genomes include both conserved regions, and hypervariable regions (HVRs), which vary highly between otherwise closely related viruses (Angly et al., 2009; Mizuno, Ghai & Rodriguez-Valera, 2014). Virally-encoded hypervariable regions can encode for host recognition structures and tail proteins (Angly et al., 2009; Garcia-Heredia et al., 2012), as well as putative internal virion proteins, terminases and ribonucleases (Mizuno, Ghai & Rodriguez-Valera, 2014; Warwick-Dugdale et al., 2019b). Previous work has shown that the hybrid, VirION approach improves capture of niche-defining HVRs (Warwick-Dugdale et al., 2019b). Our search for Sargasso Sea viral HVRs was rigorously constrained to only include regions with a minimum size ≥600 bp with near zero coverage in the top 50 most abundant viral populations to avoid false positive detection of HVRs from low-coverage taxa. Application of these parameters revealed nine putative HVRs encoded in five viral contigs; alignment of these to the NCBI NR database via a tBLASTx search (Table 1) produced functional annotations for seven of the HVRs encoded in four contigs.

**Table 1.** Predicted function of candidate hypervariable genomic regions (HVRs) in Sargasso Sea viruses obtained with the VirION pipeline. Illumina reads were mapped to the top 50 most abundant Sargasso Sea viral population representatives (at 95% identity and 70% coverage), and the encoded function of putative HVR regions that were ≥600 bp long and contained areas of zero coverage were investigated (via a tBLASTx search against the NCBI NR database).

| Contig code | HVR length | Protein name/s | Source/s |
| --- | --- | --- | --- |
| utg000082 | 993 | virion structural protein | Synechococcus phage S-CAM1 isolates<br>0910CC29; 0810SB17;<br>0809CC03; 0310NB17 |
| utg000124 | 849 | gp34; conserved hypothetical protein; **class I ribonucleotide reductase beta subunit; class I ribonucleotide reductase alpha subunit; ribonucleotide reductase domain-containing protein**; hypothetical protein ('similar to gp34'); hypothetical protein P60_gp20; hypothetical protein P60_gp21; **MazG nucleotide pyrophosphohydrolase**; | Cyanophages NATL1A-7, NATL2A-133, 9515-10a; Synechococcus phages S-CBP1, S-CBP3; Prochlorococcus phages P-SSP6, P-SSP7; Uncultured marine virus isolate Oxic3_4 |
| utg000124 | 1122 | **T7-like tail fiber** ("gp17"); major capsid protein; **phage tail fiber-like protein** | Prochlorococcus phages P-SSP7, P-SSM2; Synechococcus phage S-CBP1 |
| utg000124 | 1760 | **HNH endonuclease:HNH nuclease**; gp50; **HNH nuclease**; putative endonuclease; **phage tail fiber-like protein**; possible endonuclease; **HNH endonuclease**; HNH endonuclease family protein | Prochlorococcus phages P-SSP6, P-SSP7; P-SSP10, P-SSM2; Cyanophage 9515-10a; Prochlorococcus sp. RS04, MIT 0604; Prochlorococcus marinus str. MIT 9215; |
| utg000504 | 1986 | Hypothetical protein | **Uncultured virus clone vSAG-88-3-L14** |
| utg000504 | 3406 | **terminase small subunit; exonuclease; putative exonuclease;**<br>Hypothetical protein | Pelagibacter phage HTVC027P; Uncultured virus clone vSAG-41-H17; Pelagibacter phage HTVC023P |
| utg001170 | 933 | hypothetical protein ('**putative SAR11 Pelagibacter phage**') | Uncultured Mediterranean phage uvMED |

One advantage of long-read sequencing is the ability to identify otherwise missed HVRs – areas of viral genomes likely associated with host-virus interaction and ‘arms race’ adaptations (as proposed in host genomes; Zhao et al., 2013). Two viral contigs encoded four HVRs containing putative ORFs with homology to proteins from *Synechococcus* phages, *Prochlorococcus* phages and cyanophages, and comprised: structural proteins including T7-like tail fibre, tail fibre-like proteins, capsid proteins; MazG nucleotide pyrophosphohydrolase; ribonucleotide reductases; HNH nucleases/endonucleases (part of a homing endonuclease domain characterised by histidine and asparagine residues); and many hypothetical proteins (43% of annotations). T7-like tail fibre proteins previously recovered from pelagiphage HVRs were assumed to denote regions of rapid evolution associated with host recognition and a co-evolutionary arms race (Mizuno, Ghai & Rodriguez-Valera, 2014). This evidence suggests that phages of phototrophic hosts may also carry such genes within HVRs, supporting the paradigm of viral genome hypervariable regions as hotspots in the evolution of viral-host interactions (Avrani et al., 2011). Many HVR annotations noted here seemed tentatively unconnected to host recognition. MazG nucleotide pyrophosphohydrolase may function in phage DNA synthesis. A previously hypothesised role in phage mediated regulation of the stringent response (i.e., host reaction to nutrient deprivation; Bryan et al., 2008), has recently been revised, and MazG is now thought to enable recycling of host DNA via the hydrolysis of deoxyribonucleotides (Rihtman et al., 2019). Enzymes such as ribonucleotide reductase (RNR), and HNH nucleases and endonucleases are predicted to function in phage DNA synthesis and packaging (respectively). RNR is a viral AMG that catalyses the formation of deoxyribonucleotides from ribonucleotides for the production of progeny DNA (Nordlund & Reichard, 2006; Thompson et al., 2011). RNR genes are abundant and under selective evolutionary pressure in environmental viral assemblages (Dwivedi et al., 2013). HNH proteins function in conjunction with terminase in DNA cleavage for phage morphogenesis and are widespread in long-tailed phages (Kala et al., 2014). HNH nucleases have been identified in phages from disparate environments, including a deep-sea thermophilic bacteriophage (Zhang et al., 2017) and ‘hidden’ prophage of a cultured marine *Roseobacter* (Zhao et al., 2010).

Why genes for DNA synthesis and packaging enzymes, and even genes related to viral structure (i.e., encoding virion and capsid proteins) have been observed within HVRs may reflect an aspect of phage biology not yet understood (Mizuno, Ghai & Rodriguez-Valera, 2014). However, the detection of HNH nucleases here could suggest that the HVRs of viruses infecting phototrophs encode proteins to counter anti-phage defences beyond adsorption and injection of viral DNA. A potential mechanism for overcoming host defence is concerned with avoidance of Restriction-Modification (RM) systems which have been identified in multiple strains of *Prochlorococcus*, but are rare in SAR11 (Giovannoni, 2017; Chen et al., 2019). RM systems identify and destroy viral DNA: phages can evade detection by the removal, under-representation or mutation of the restriction-site sequences recognised by the host cell (Kruger & Bickle, 1983; Hampton, Watson & Fineran, 2020). Phage encoded HNH type nucleases have been identified as likely protagonists in viral genomic rearrangement (Hatfull, 2014; *from Life in Phage World: 5-57*), so have the potential to assist phage avoidance of host RM systems via this route.

The remaining three HVRs encoded by two contigs may have derived from pelagiphages. Exonuclease, terminase and hypothetical protein annotations were sourced from pelagiphages HTVC027P and HTV023P, a putative pelagiphage acquired through metagenomic fosmids (from the Mediterranean deep chlorophyll maximum; (Mizuno et al., 2013), and viral genomes recovered from single-virus genomics, including a virus sampled from the bathypelagic (‘vSAG-88-3-L14’), and another described as one of the most abundant dsDNA viruses in the surface global marine virosphere (at species level; ‘vSAG-41-H17’) (Martinez-Hernandez et al., 2017). Like HNH proteins, exonucleases and terminases are predicted to function in phage DNA packaging (Son & Serwer, 1992; Kala et al., 2014), and have been previously identified in pelagiphage genomes (Zhao, Qin & Zhang, 2019; Buchholz et al., 2021b), and viral metagenomic HVRs (Mizuno, Ghai & Rodriguez-Valera, 2014; Warwick-Dugdale et al., 2019b). However, it is worth noting that the terminase located in the HVR did not cluster with terminases from known pelagiphages in a phylogenetic tree (Figure S6) and it is unknown as to whether HVR structure provides a robust signal for host association.

## CONCLUSIONS

Although host prediction through bioinformatic approaches proved challenging, our results appear to support previous evidence that SAR11 viruses are not abundant in Sargasso Sea communities (Parsons et al., 2012). Surprisingly, our results also suggest that the viruses of cyanobacteria are not abundant in the Sargasso Sea. In the context of warming global oceans, the possibility that oligotrophic environments such as the Sargasso Sea have lower rates of active lytic viral replication in the dominant members and lower viral macrodiversity poses important questions for understanding marine carbon cycling. SAR11 are the most abundant bacteria in the oceans, but are most dominant in stratified, nutrient-poor regions of the ocean (Giovannoni, 2017). As top-down controllers of microbial communities, viruses are a major constituent of global carbon cycles. SAR11 consume the organic carbon and release CO_2_, but simultaneously, viral mediated lysis works to return cells to dissolved organic matter (DOM), available then for microbial consumption (the ‘Viral Shunt’; Wilhelm & Suttle, 1999). If stratified, nutrient-poor regions are generally characterised not only by highly abundant SAR11, but also low rates of viral-mediated host lysis, there are implications for the way in which carbon will flow through increasingly vast areas of the future global ocean.

## ACKNOWLEDGEMENTS

The authors thank the crew and the marine technicians of the Bermuda Institute of Ocean Science vessel, the ‘Atlantic Explorer’, for the collection of seawater samples. Bioinformatic analyses were conducted using the high-performance computing resources of the Ohio Supercomputer Center (OSC, 1987), provided by the Louisiana State University and those of ISCA, provided by the University of Exeter.

## ADDITIONAL INFORMATION AND DECLARATIONS

### Author Contributions

- Joanna Warwick-Dugdale^**†**^ performed the experiments, analyzed the data, prepared figures and/or tables, authored or reviewed drafts of the paper, approved the final draft.
- Funing Tian^**†**^ analyzed the data, prepared figures and/or tables, authored or reviewed drafts of the paper, approved the final draft.
- Michelle Michelsen performed the experiments.
- Dylan R Cronin analyzed the data.
- Karen Moore performed the experiments, contributed reagents/materials/analysis tools.
- Audrey Farbos performed the experiments.
- Lauren Chittick performed the experiments.
- Ashley Bell analyzed the data and prepared figures.
- Holger Buchholz analyzed the data.
- Rachel Parsons contributed reagents/materials/analysis tools, authored or reviewed drafts of the paper, approved the final draft
- Ahmed A Zayed analyzed the data.
- Michael J Allen authored or reviewed drafts of the paper, approved the final draft
- Matthew B Sullivan contributed reagents/materials/analysis tools, authored or reviewed drafts of the paper, approved the final draft.
- Ben Temperton conceived and designed the experiments, analyzed the data, contributed reagents/materials/analysis tools, prepared figures and/or tables, authored or reviewed drafts of the paper, approved the final draft.

^**†**^The first two authors should be regarded as Joint First Authors.

### DNA Deposition

Sequencing data and assemblies are available at the National Center for Biotechnology Information under the BioProject accession number PRJNA767318

### Data Availability

All code and analyses can be found at: https://www.protocols.io/private/8E071162E5D611ECB22B0A58A9FEAC0

### Supplemental Information

Supplemental information for this article can be found online at

## REFERENCES

Alonso-Sáez L, Morán XAG, Clokie MR. 2018. Low activity of lytic pelagiphages in coastal marine waters. ISME Journal 12:2100–2102. DOI: 10.1038/s41396-018-0185-y.

Angly FE, Felts B, Breitbart M, Salamon P, Edwards RA, Carlson C, Chan AM, Haynes M, Kelley S, Liu H, Mahaffy JM, Mueller JE, Nulton J, Olson R, Parsons R, Rayhawk S, Suttle CA, Rohwer F. 2006. The marine viromes of four oceanic regions. PLoS Biology 4:2121–2131. DOI: 10.1371/journal.pbio.0040368.

Angly F, Youle M, Nosrat B, Srinagesh S, Rodriguez-Brito B, McNairnie P, Deyanat-Yazdi G, Breitbart M, Rohwer F. 2009. Genomic analysis of multiple Roseophage SIO1 strains. Environmental Microbiology 11:2863–2873. DOI: 10.1111/j.1462-2920.2009.02021.x.

Avrani S, Wurtzel O, Sharon I, Sorek R, Lindell D. 2011. Genomic island variability facilitates Prochlorococcus-virus coexistence. Nature 474:604–608. DOI: 10.1038/nature10172.

Baxter JM. 2016. Explaining Ocean Warming: Causes, scale, effects and consequences. IUCN, International Union for Conservation of Nature. DOI: 10.2305/IUCN.CH.2016.08.en.

Breitbart M, Bonnain C, Malki K, Sawaya NA. 2018. Phage puppet masters of the marine microbial realm. Nature Microbiology 3. DOI: 10.1038/s41564-018-0166-y.

Brockhurst MA, Buckling A, Rainey PB. 2005. The effect of a bacteriophage on diversification of the opportunistic bacterial pathogen, Pseudomonas aeruginosa. Proceedings of the Royal Society B: Biological Sciences 272:1385–1391. DOI: 10.1098/rspb.2005.3086.

Brum JR, Ignacio-espinoza JC, Roux S, Doulcier G, Acinas SG, Alberti A, Chaffron S. 2015. Patterns and ecological drivers of ocean viral communities. Science 348:1261498-1–11. DOI: 10.1126/science.1261498.

Bryan MJ, Burroughs NJ, Spence EM, Clokie MRJ, Mann NH, Bryan SJ. 2008. Evidence for the intense exchange of MazG in marine cyanophages by horizontal gene transfer. PLoS ONE 3:1–12. DOI: 10.1371/journal.pone.0002048.

Buchholz HH, Bolaños LM, Bell AG, Michelsen ML, Allen MJ. 2021a. Genomic evidence for inter-class host transition between abundant streamlined heterotrophs by a novel and ubiquitous marine Methylophage. bioRxiv. DOI: 10.1101/2021.08.24.457595.

Buchholz HH, Michelsen ML, Bolaños LM, Browne E, Allen MJ, Temperton B. 2021b. Efficient dilution-to-extinction isolation of novel virus–host model systems for fastidious heterotrophic bacteria. ISME Journal. DOI: 10.1038/s41396-020-00872-z.

Calcagno V, Jarne P, Loreau M, Mouquet N, David P. 2017. Diversity spurs diversification in ecological communities. Nature Communications 8:1–9. DOI: 10.1038/ncomms15810.

Capella-Gutiérrez S, Silla-Martínez JM, Gabaldón T. 2009. trimAl: A tool for automated alignment trimming in large-scale phylogenetic analyses. Bioinformatics 25:1972–1973. DOI: 10.1093/bioinformatics/btp348.

Capotondi A, Alexander MA, Bond NA, Curchitser EN, Scott JD. 2012. Enhanced upper ocean stratification with climate change in the CMIP3 models. Journal of Geophysical Research: Oceans 117:1–23. DOI: 10.1029/2011JC007409.

Carlson CA, Morris R, Parsons R, Treusch AH, Giovannoni SJ, Vergin K. 2009. Seasonal dynamics of SAR11 populations in the euphotic and mesopelagic zones of the northwestern Sargasso Sea. ISME Journal 3:283–295. DOI: 10.1038/ismej.2008.117.

Chen L-X, Zhao Y, McMahon KD, Mori JF, Jessen GL, Nelson TC, Warren LA, Banfield JF. 2019. Wide Distribution of Phage That Infect Freshwater SAR11 Bacteria. mSystems 4:1–16. DOI: 10.1128/msystems.00410-19.

Coleman ML, Sullivan MB, Martiny AC, Steglich C, Barry K, DeLong EF, Chisholm SW. 2006. Genomic islands and the ecology and evolution of Prochlorococcus. Science 311:1768–1770. DOI: 10.1126/science.1122050.

Cornillon P, Stramma L. 1985. The distribution of diurnal sea surface warming events in the western Sargasso Sea. Journal of Geophysical Research 90:11811. DOI: 10.1029/JC090iC06p11811.

De Coster W, D’Hert S, Schultz DT, Cruts M, Van Broeckhoven C. 2018. NanoPack: Visualizing and processing long-read sequencing data. Bioinformatics 34:2666–2669. DOI: 10.1093/bioinformatics/bty149.

Cottrell MT, Kirchman DL. 2016. Transcriptional control in marine copiotrophic and oligotrophic bacteria with streamlined genomes. Applied and Environmental Microbiology 82:6010–6018. DOI: 10.1128/AEM.01299-16.

Cumming G. 2014. The New Statistics : Why and How. Phychological Science 25:7–29. DOI: 10.1177/0956797613504966.

Dwivedi B, Xue B, Lundin D, Edwards RA, Breitbart M. 2013. A bioinformatic analysis of ribonucleotide reductase genes in phage genomes and metagenomes. BMC Evolutionary Biology 13:1–17. DOI: 10.1186/1471-2148-13-33.

Emerson JB, Roux S, Brum JR, Bolduc B, Woodcroft BJ, Jang H Bin, Singleton CM, Solden LM, Naas AE, Boyd JA, Hodgkins SB, Wilson RM, Trubl G, Li C, Frolking S, Pope PB, Wrighton KC, Crill PM, Chanton JP, Saleska SR, Tyson GW, Rich VI, Sullivan MB. 2018. Host-linked soil viral ecology along a permafrost thaw gradient. Nature Microbiology 3:870–880. DOI: 10.1038/s41564-018-0190-y.

Falkowski PG, Fenchel T, Delong EF. 2008. The Microbial Engines That Drive Earth’s Biogeochemical Cycles. Science 320:1034–1039. DOI: 10.1126/science.1153213.

Forterre P. 2012. The virocell concept and environmental microbiology. The ISME Journal 7:233–236. DOI: 10.1038/ismej.2012.110.

Fuhrman JA, Comeau DE, Hagström A, Chan AM. 1988. Extraction from natural planktonic microorganisms of DNA suitable for molecular biological studies. Applied and Environmental Microbiology 54:1426–9.

Galiez C, Siebert M, Enault F, Vincent J, Söding J. 2017. WIsH: who is the host? Predicting prokaryotic hosts from metagenomic phage contigs. Bioinformatics 33:3113–3114. DOI: 10.1093/bioinformatics/btx383.

Garcia-Heredia I, Martin-Cuadrado AB, Mojica FJM, Santos F, Mira A, Antón J, Rodriguez-Valera F. 2012. Reconstructing viral genomes from the environment using fosmid clones: The case of haloviruses. PLoS ONE 7. DOI: 10.1371/journal.pone.0033802.

Giovannoni SJ. 1990. Genetic diversity in Sargasso sea bacterioplankton. Nature 345:183–187. DOI: 10.1038/346183a0.

Giovannoni SJ. 2017. SAR11 Bacteria: The Most Abundant Plankton in the Oceans. Annual Review of Marine Science 9:231–255. DOI: 10.1146/annurev-marine-010814-015934.

Giovannoni SJ, DeLong EF, Schmidt TM, Pace NR. 1990. Tangential flow filtration and preliminary phylogenetic analysis of marine picoplankton. Applied and Environmental Microbiology 56:2572–2575.

Giovannoni SJ, Rappé MS, Vergin KL, Adair NL. 1996. 16S rRNA genes reveal stratified open ocean bacterioplankton populations related to the green nonsulfur bacteria. Proceedings of the National Academy of Sciences of the United States of America 93:7979–7984. DOI: 10.1073/pnas.93.15.7979.

Giovannoni SJ, Vergin KL. 2012. Seasonality in ocean microbial communities. Science 335:671–676. DOI: 10.1126/science.1198078.

Gordon DA, Giovannoni SJ. 1996. Detection of stratified microbial populations related to Chlorobium and Fibrobacter species in the Atlantic and Pacific Oceans. Applied and Environmental Microbiology 62:1171–1177. DOI: 10.1128/aem.62.4.1171-1177.1996.

Gregory AC, Gerhardt K, Zhong Z-P, Bolduc B, Temperton B, Konstantinidis KT, Sullivan MB. 2020a. MetaPop: A pipeline for macro-and micro-diversity analyses and visualization of microbial and viral metagenome-derived populations. bioRxiv. DOI: https://doi.org/10.1101/2020.11.01.363960.

Gregory AC, Zablocki O, Zayed AA, Howell A, Bolduc B, Sullivan MB. 2020b. The Gut Virome Database Reveals Age-Dependent Patterns of Virome Diversity in the Human Gut. Cell Host and Microbe 28:724–740.e8. DOI: 10.1016/j.chom.2020.08.003.

Gregory AC, Zayed AA, Conceição-Neto N, Temperton B, Bolduc B, Alberti A, Ardyna M, Arkhipova K, Carmichael M, Cruaud C, Dimier C, Domínguez-Huerta G, Ferland J, Kandels S, Liu Y, Marec C, Pesant S, Picheral M, Pisarev S, Poulain J, Tremblay JÉ, Vik D, Acinas SG, Babin M, Bork P, Boss E, Bowler C, Cochrane G, de Vargas C, Follows M, Gorsky G, Grimsley N, Guidi L, Hingamp P, Iudicone D, Jaillon O, Kandels-Lewis S, Karp-Boss L, Karsenti E, Not F, Ogata H, Poulton N, Raes J, Sardet C, Speich S, Stemmann L, Sullivan MB, Sunagawa S, Wincker P, Culley AI, Dutilh BE, Roux S. 2019. Marine DNA Viral Macro-and Microdiversity from Pole to Pole. Cell 177:1109–1123.e14. DOI: 10.1016/j.cell.2019.03.040.

Guidi L, Chaffron S, Bittner L, Eveillard D, Larhlimi A, Roux S, Darzi Y, Audic S, Berline L, Brum JR, Coelho LP, Espinoza JCI, Malviya S, Sunagawa S, Dimier C, Kandels-Lewis S, Picheral M, Poulain J, Searson S, Stemmann L, Not F, Hingamp P, Speich S, Follows M, Karp-Boss L, Boss E, Ogata H, Pesant S, Weissenbach J, Wincker P, Acinas SG, Bork P, De Vargas C, Iudicone D, Sullivan MB, Raes J, Karsenti E, Bowler C, Gorsky G. 2016. Plankton networks driving carbon export in the oligotrophic ocean. Nature 532:465–470. DOI: 10.1038/nature16942.

Guo J, Bolduc B, Zayed AA, Varsani A, Dominguez-Huerta G, Delmont TO, Pratama AA, Gazitúa MC, Vik D, Sullivan MB, Roux S. 2021. VirSorter2: a multi-classifier, expert-guided approach to detect diverse DNA and RNA viruses. Microbiome 9:1–13. DOI: 10.1186/s40168-020-00990-y.

Hampton HG, Watson BNJ, Fineran PC. 2020. The arms race between bacteria and their phage foes. Nature 577:327–336. DOI: 10.1038/s41586-019-1894-8.

Hart SP, Schreiber SJ, Levine JM. 2016. How variation between individuals affects species coexistence. Ecology letters 19:825–838. DOI: 10.1111/ele.12618.

Hatfull GF. 2014. Unintelligent Design: Mosaic Construction of Bacteriophage Genomes. In: Rohwer FL, Youle M, Maughan H, Hisakawa N eds. Life in Our Phage World: A centennial field guide to the Earth’s most diverse inhabitants. San Diego, CA: Wholon, 5–47.

Henson MW, Lanclos VC, Pitre DM, Weckhorst JL, Lucchesi AM, Cheng C, Temperton B, Thrash JC. 2020. Expanding the diversity of bacterioplankton isolates and modeling isolation efficacy with large scale dilution-to-extinction cultivation. Applied and environmental microbiology. DOI: 10.1128/AEM.00943-20.

Howard-Varona C, Lindback MM, Bastien GE, Solonenko N, Zayed AA, Jang H bin, Andreopoulos B, Brewer HM, Glavina del Rio T, Adkins JN, Paul S, Sullivan MB, Duhaime MB. 2020. Phage-specific metabolic reprogramming of virocells. ISME Journal 14:881–895. DOI: 10.1038/s41396-019-0580-z.

Hyatt D, Chen G-L, LoCascio PF, Land ML, Larimer FW, Hauser LJ. 2010. Prodigal: prokaryotic gene recognition and translation initiation site identification. BMC Bioinformatics 11:119. DOI: 10.1186/1471-2105-11-119.

Jacquet S, Partensky F, Lennon JF, Vaulot D. 2001. Diel patterns of growth and division in marine picoplankton in culture. Journal of Phycology 37:357–369. DOI: 10.1046/j.1529-8817.2001.037003357.x.

Bin Jang H, Bolduc B, Zablocki O, Kuhn JH, Roux S, Adriaenssens EM, Brister JR, Kropinski AM, Krupovic M, Lavigne R, Turner D, Sullivan MB. 2019. Taxonomic assignment of uncultivated prokaryotic virus genomes is enabled by gene-sharing networks. Nature Biotechnology 37:632–639. DOI: 10.1038/s41587-019-0100-8.

John SG, Mendez CB, Deng L, Poulos B, Kauffman AKM, Kern S, Brum J, Polz MF, Boyle E a, Sullivan MB. 2011. A simple and efficient method for concentration of ocean viruses by chemical flocculation. Environmental microbiology reports 3:195–202. DOI: 10.1111/j.1758-2229.2010.00208.x.

Kala S, Cumby N, Sadowski PD, Hyder BZ, Kanelis V, Davidson AR, Maxwell KL. 2014. HNH proteins are a widespread component of phage DNA packaging machines. Proceedings of the National Academy of Sciences of the United States of America 111:6022–6027. DOI: 10.1073/pnas.1320952111.

Kalyaanamoorthy S, Minh BQ, Wong TKF, Von Haeseler A, Jermiin LS. 2017. ModelFinder: Fast model selection for accurate phylogenetic estimates. Nature Methods 14:587–589. DOI: 10.1038/nmeth.4285.

Kang I, Oh H-M, Kang D, Cho J-C. 2013. Genome of a SAR116 bacteriophage shows the prevalence of this phage type in the oceans. Proceedings of the National Academy of Sciences of the United States of America 110:12343–8. DOI: 10.1073/pnas.1219930110.

Kashtan N, Roggensack SE, Rodrigue S, Thompson JW, Biller SJ, Coe A, Ding H, Marttinen P, Malmstrom RR, Stocker R, Follows MJ, Stepanauskas R, Chisholm SW. 2014. Single-cell genomics reveals hundreds of coexisting subpopulations in wild Prochlorococcus. Science 344:416–420. DOI: 10.1126/science.1248575.

Katoh K, Standley DM. 2016. A simple method to control over-alignment in the MAFFT multiple sequence alignment program. Bioinformatics 32:1933–1942. DOI: 10.1093/bioinformatics/btw108.

Kearse M, Moir R, Wilson A, Stones-Havas S, Cheung M, Sturrock S, Buxton S, Cooper A, Markowitz S, Duran C, Thierer T, Ashton B, Meintjes P, Drummond A. 2012. Geneious Basic: An integrated and extendable desktop software platform for the organization and analysis of sequence data. Bioinformatics 28:1647–1649. DOI: 10.1093/bioinformatics/bts199.

Kelly L, Ding H, Huang KH, Osburne MS, Chisholm SW. 2013. Genetic diversity in cultured and wild marine cyanomyoviruses reveals phosphorus stress as a strong selective agent. ISME Journal 7:1827–1841. DOI: 10.1038/ismej.2013.58.

Khot V, Strous M, Hawley AK. 2020. Computational approaches in viral ecology. Computational and Structural Biotechnology Journal 18:1605–1612. DOI: 10.1016/j.csbj.2020.06.019.

Kolmogorov M, Bickhart DM, Behsaz B, Gurevich A, Rayko M, Shin SB, Kuhn K, Yuan J, Polevikov E, Smith TPL, Pevzner PA. 2020. metaFlye: scalable long-read metagenome assembly using repeat graphs. Nature Methods 17:1103–1110. DOI: 10.1038/s41592-020-00971-x.

Krishnamurthy SR, Wang D. 2017. Origins and challenges of viral dark matter. Virus Research 239:136–142. DOI: 10.1016/j.virusres.2017.02.002.

Kruger DH, Bickle TA. 1983. Bacteriophage survival: Multiple mechanisms for avoiding the deoxyribonucleic acid restriction systems of their hosts. Microbiological Reviews 47:345–360. DOI: 10.1128/mmbr.47.3.345-360.1983.

Langmead B, Salzberg SL. 2012. Fast gapped-read alignment with Bowtie 2. Nature Methods 9:357–359. DOI: 10.1038/nmeth.1923.

Letunic I, Bork P. 2016. Interactive tree of life (iTOL) v3: an online tool for the display and annotation of phylogenetic and other trees. Nucleic acids research 44:W242–W245. DOI: 10.1093/nar/gkw290.

Lindell D, Jaffe JD, Coleman ML, Futschik ME, Axmann IM, Rector T, Kettler G, Sullivan MB, Steen R, Hess WR, Church GM, Chisholm SW. 2007. Genome-wide expression dynamics of a marine virus and host reveal features of coevolution. Nature 449:83–86. DOI: 10.1038/nature06130.

Lindell D, Sullivan MB, Johnson ZI, Tolonen AC, Rohwer F, Chisholm SW. 2004. Transfer of photosynthesis genes to and from Prochlorococcus viruses. Proceedings of the National Academy of Sciences of the United States of America 101:11013–11018. DOI: 10.1073/pnas.0401526101.

Lomas MW, Bates NR, Buck KN, Knap AH. 2011. Oceanography of the Sargasso Sea: Overview of Scientific Studies.

Lomas MW, Bates NR, Johnson RJ, Knap AH, Steinberg DK, Carlson CA. 2013. Two decades and counting: 24-years of sustained open ocean biogeochemical measurements in the Sargasso Sea. Deep Sea Research Part II: Topical Studies in Oceanography 93:16–32. DOI: 10.1016/j.dsr2.2013.01.008.

López-Pérez M, Haro-Moreno JM, Coutinho FH, Martinez-Garcia M, Rodriguez-Valera F. 2020. The Evolutionary Success of the Marine Bacterium SAR11 Analyzed through a Metagenomic Perspective. mSystems 5:1–13. DOI: 10.1128/msystems.00605-20.

Madi N, Vos M, Murall CL, Legendre P, Shapiro BJ. 2020. Does diversity beget diversity in microbiomes? eLife 9:1–83. DOI: 10.7554/eLife.58999.

Martin M. 2011. Cutadapt removes adapter sequences from high-throughput sequencing reads. EMBnet.journal 17:10. DOI: 10.14806/ej.17.1.200.

Martinez-Hernandez F, Fornas O, Lluesma Gomez M, Bolduc B, de la Cruz Peña MJ, Martínez JM, Anton J, Gasol JM, Rosselli R, Rodriguez-Valera F, Sullivan MB, Acinas SG, Martinez-Garcia M. 2017. Single-virus genomics reveals hidden cosmopolitan and abundant viruses. Nature communications 8:15892. DOI: 10.1038/ncomms15892.

Martinez-Hernandez F, Fornas Ò, Lluesma Gomez M, Garcia-Heredia I, Maestre-Carballa L, López-Pérez M, Haro-Moreno JM, Rodriguez-Valera F, Martinez-Garcia M. 2019. Single-cell genomics uncover Pelagibacter as the putative host of the extremely abundant uncultured 37-F6 viral population in the ocean. ISME Journal 13:232–236. DOI: 10.1038/s41396-018-0278-7.

Menzel DW, Ryther JH. 1959. The annual cycle of primary production in the Sargasso Sea off Bermuda. Deep Sea Research (1953) 6:351–367. DOI: 10.1016/0146-6313(59)90095-4.

Mizuno CM, Ghai R, Rodriguez-Valera F. 2014. Evidence for metaviromic islands in marine phages. Frontiers in Microbiology 5:1–10. DOI: 10.3389/fmicb.2014.00027.

Mizuno CM, Rodriguez-Valera F, Kimes NE, Ghai R. 2013. Expanding the Marine Virosphere Using Metagenomics. PLoS Genetics 9:e1003987. DOI: 10.1371/journal.pgen.1003987.

Moon K, Cho J. 2021. Metaviromics coupled with phage-host identification to open the viral ‘black box.’ Journal of Microbiology 59:311–323. DOI: 10.1007/s12275-021-1016-9.

Moore L, Rocap G. 1998. Physiology and molecular phylogeny of coexisting Prochlorococcus ecotypes. Nature 576:220–223.

Morris RM, Rappé MS, Connon S a, Vergin KL, Siebold W a, Carlson C a, Giovannoni SJ. 2002. SAR11 clade dominates ocean surface bacterioplankton communities. Nature 420:806–810. DOI: 10.1038/nature01240.

Morris RM, Vergin KL, Cho JC, Rappé MS, Carlson CA, Giovannoni SJ. 2005. Temporal and spatial response of bacterioplankton lineages to annual convective overturn at the Bermuda Atlantic Time-series Study site. Limnology and Oceanography 50:1687–1696. DOI: 10.4319/lo.2005.50.5.1687.

Mruwat N, Carlson MCG, Goldin S, Ribalet F, Kirzner S, Hulata Y, Beckett SJ, Shitrit D, Weitz JS, Armbrust EV, Lindell D. 2020. A single-cell polony method reveals low levels of infected Prochlorococcus in oligotrophic waters despite high cyanophage abundances. ISME Journal. DOI: 10.1038/s41396-020-00752-6.

Nayfach S, Camargo AP, Schulz F, Eloe-Fadrosh E, Roux S, Kyrpides NC. 2020. CheckV assesses the quality and completeness of metagenome-assembled viral genomes. Nature Biotechnology. DOI: 10.1038/s41587-020-00774-7.

Nei M, Li WH. 1979. Mathematical model for studying genetic variation in terms of restriction endonucleases. Proceedings of the National Academy of Sciences of the United States of America 76:5269–5273. DOI: 10.1073/pnas.76.10.5269.

Nguyen LT, Schmidt HA, Von Haeseler A, Minh BQ. 2015. IQ-TREE: A fast and effective stochastic algorithm for estimating maximum-likelihood phylogenies. Molecular Biology and Evolution 32:268–274. DOI: 10.1093/molbev/msu300.

Noble RT, Fuhrman JA. 2000. Rapid virus production and removal as measured with fluorescently labeled viruses as tracers. Applied and Environmental Microbiology 66:3790–3797. DOI: 10.1128/AEM.66.9.3790-3797.2000.

Nordlund P, Reichard P. 2006. Ribonucleotide reductases. Annual Review of Biochemistry 75:681–706. DOI: 10.1146/annurev.biochem.75.103004.142443.

Nurk S, Meleshko D, Korobeynikov A, Pevzner PA. 2017. MetaSPAdes: A new versatile metagenomic assembler. Genome Research 27:824–834. DOI: 10.1101/gr.213959.116.

Oksanen J, Blanchet FG, Friendly M, Kindt R, Legendre P, Mcglinn D, Minchin PR, O’hara RB, Simpson GL, Solymos P, Henry M, Stevens H, Szoecs E, Maintainer HW. 2020. Package “vegan” Title Community Ecology Package Version 2.5–7.

Ottesen EA, Young CR, Gifford SM, Eppley JM, Marin R, Schuster SC, Scholin CA, DeLong. E. F. 2014. Multispecies diel transcriptional oscillations in open ocean heterotrophic bacterial assemblages Accessed Title : Multispecies d iel transcriptional oscillations in open ocean. Science 345:207–212.

Paez-Espino D, Eloe-Fadrosh EA, Pavlopoulos GA, Thomas AD, Huntemann M, Mikhailova N, Rubin E, Ivanova NN, Kyrpides NC. 2016. Uncovering Earth’s virome. Nature 536:425–430. DOI: 10.1038/nature19094.

Parsons RJ, Breitbart M, Lomas MW, Carlson CA. 2012. Ocean time-series reveals recurring seasonal patterns of virioplankton dynamics in the northwestern Sargasso Sea. The ISME Journal 6:273–284. DOI: 10.1038/ismej.2011.101.

Rihtman B, Bowman-Grahl S, Millard A, Corrigan RM, Clokie MRJ, Scanlan DJ. 2019. Cyanophage MazG is a pyrophosphohydrolase but unable to hydrolyse magic spot nucleotides. Environmental Microbiology Reports 11:448–455. DOI: 10.1111/1758-2229.12741.

Rodriguez-Valera F, Martin-Cuadrado A-B, Rodriguez-Brito B, Pasić L, Thingstad TF, Rohwer F, Mira A. 2009. Explaining microbial population genomics through phage predation. Nature reviews. Microbiology 7:828–36. DOI: 10.1038/nrmicro2235.

Roux S, Emerson JB, Eloe-Fadrosh EA, Sullivan MB. 2017. Benchmarking viromics: An in silico evaluation of metagenome-enabled estimates of viral community composition and diversity. PeerJ:1–26. DOI: 10.7287/PEERJ.PREPRINTS.3053V1.

Roux S, Enault F, Hurwitz BL, Sullivan MB. 2015a. VirSorter: mining viral signal from microbial genomic data. PeerJ 3:e985. DOI: 10.7717/peerj.985.

Roux S, Hallam SJ, Woyke T, Sullivan MB. 2015b. Viral dark matter and virus – host interactions resolved from publicly available microbial genomes. eLife 4:1–20. DOI: 10.7554/eLife.08490.

Rusch DB, Halpern AL, Sutton G, Heidelberg KB, Williamson S, Yooseph S, Wu D, Eisen JA, Hoffman JM, Remington K, Beeson K, Tran B, Smith H, Baden-Tillson H, Stewart C, Thorpe J, Freeman J, Andrews-Pfannkoch C, Venter JE, Li K, Kravitz S, Heidelberg JF, Utterback T, Rogers YH, Falcón LI, Souza V, Bonilla-Rosso G, Eguiarte LE, Karl DM, Sathyendranath S, Platt T, Bermingham E, Gallardo V, Tamayo-Castillo G, Ferrari MR, Strausberg RL, Nealson K, Friedman R, Frazier M, Venter JC. 2007. The Sorcerer II Global Ocean Sampling expedition: Northwest Atlantic through eastern tropical Pacific. PLoS Biology 5:0398–0431. DOI: 10.1371/journal.pbio.0050077.

Santos-Medellin C, Zinke LA, ter Horst AM, Gelardi DL, Parikh SJ, Emerson JB. 2021. Viromes outperform total metagenomes in revealing the spatiotemporal patterns of agricultural soil viral communities. ISME Journal. DOI: 10.1038/s41396-021-00897-y.

Scanlan PD, Hall AR, Blackshields G, Friman V-P, Davis MR, Goldberg JB, Buckling A. 2015. Coevolution with bacteriophages drives genome-wide host evolution and constrains the acquisition of abiotic-beneficial mutations. Molecular biology and evolution 32:1425–35. DOI: 10.1093/molbev/msv032.

Schroeder PJ, Jenkins DG. 2018. How robust are popular beta diversity indices to sampling error. Ecosphere 9. DOI: 10.1002/ecs2.2100.

Shaffer M, Borton MA, McGivern BB, Zayed AA, La Rosa SL 0003 3527 8101, Solden LM, Liu P, Narrowe AB, Rodríguez-Ramos J, Bolduc B, Gazitúa MC, Daly RA, Smith GJ, Vik DR, Pope PB, Sullivan MB, Roux S, Wrighton KC. 2020. DRAM for distilling microbial metabolism to automate the curation of microbiome function. Nucleic Acids Research 48:8883–8900. DOI: 10.1093/nar/gkaa621.

Smyth TJ, Allen I, Atkinson A, Bruun JT, Harmer RA, Pingree RD, Widdicombe CE, Somerfield PJ. 2014. Ocean net heat flux influences seasonal to interannual patterns of plankton abundance. PLoS ONE 9. DOI: 10.1371/journal.pone.0098709.

Son M, Serwer P. 1992. Role of Exonuclease in the Specificity of Bacteriophage T7 DNA Packaging. Virology 190:824–833.

Steinberg DK, Carlson CA, Bates NR, Johnson RJ, Michaels AF, Knap AH. 2001. Overview of the US JGOFS Bermuda Atlantic Time-series Study (BATS): A decade-scale look at ocean biology and biogeochemistry. Deep-Sea Research Part II: Topical Studies in Oceanography 48:1405–1447. DOI: 10.1016/S0967-0645(00)00148-X.

Stewart RD, Auffret MD, Warr A, Walker AW, Roehe R, Watson M. 2019. Compendium of 4,941 rumen metagenome-assembled genomes for rumen microbiome biology and enzyme discovery. Nature Biotechnology 37:953–961. DOI: 10.1038/s41587-019-0202-3.

Sullivan MB, Coleman ML, Weigele P, Rohwer F, Chisholm SW. 2005. Three Prochlorococcus cyanophage genomes: Signature features and ecological interpretations. PLoS Biology 3:0790–0806. DOI: 10.1371/journal.pbio.0030144.

Sullivan MB, Huang KH, Ignacio-Espinoza JC, Berlin AM, Kelly L, Weigele PR, DeFrancesco AS, Kern SE, Thompson LR, Young S, Yandava C, Fu R, Krastins B, Chase M, Sarracino D, Osburne MS, Henn MR, Chisholm SW. 2010. Genomic analysis of oceanic cyanobacterial myoviruses compared with T4-like myoviruses from diverse hosts and environments. Environmental Microbiology 12:3035–3056. DOI: 10.1111/j.1462-2920.2010.02280.x.

Sullivan MB, Lindell D, Lee JA, Thompson LR, Bielawski JP, Chisholm SW. 2006. Prevalence and evolution of core photosystem II genes in marine cyanobacterial viruses and their hosts. PLoS Biology 4:1344–1357. DOI: 10.1371/journal.pbio.0040234.

Suttle CA. 2007. Marine viruses--major players in the global ecosystem. Nature reviews. Microbiology 5:801–812. DOI: 10.1038/nrmicro1750.

Suttle CA, Chen F. 1992. Mechanisms and rates of decay of marine viruses in seawater. Applied and Environmental Microbiology 58:3721–3729. DOI: 10.1128/aem.58.11.3721-3729.1992.

Thompson LR, Zeng Q, Kelly L, Huang KH, Singer AU, Stubbe J, Chisholm SW. 2011. Phage auxiliary metabolic genes and the redirection of cyanobacterial host carbon metabolism. Proceedings of the National Academy of Sciences 108:E757–E764. DOI: 10.1073/pnas.1102164108.

Treusch AH, Vergin KL, Finlay LA, Donatz MG, Burton RM, Carlson CA, Giovannoni SJ. 2009. Seasonality and vertical structure of microbial communities in an ocean gyre. The ISME Journal 3:1148–1163. DOI: 10.1038/ismej.2009.60.

Trubl G, Jang H Bin, Roux S, Emerson JB, Solonenko N, Vik DR, Solden L, Ellenbogen J, Runyon AT, Bolduc B, Woodcroft BJ, Saleska SR, Tyson GW, Wrighton KC, Sullivan MB, Rich VI. 2018. Soil Viruses Are Underexplored Players in Ecosystem Carbon Processing. mSystems 3:1–21. DOI: 10.1128/msystems.00076-18.

Vaser R, Sović I, Nagarajan N, Šikić M. 2017. Fast and accurate de novo genome assembly from long uncorrected reads. Genome Research 27:737–746. DOI: 10.1101/gr.214270.116.

Venter JC, Remington K, Heidelberg JF, Halpern AL, Rusch D, Eisen JA, Wu D, Paulsen I, Nelson KE, Nelson W, Fouts DE, Levy S, Knap AH, Lomas MW, Nealson K, White O, Peterson J, Hoffman J, Parsons R, Baden-Tillson H, Pfannkoch C, Rogers YH, Smith HO. 2004. Environmental Genome Shotgun Sequencing of the Sargasso Sea. Science 304:66–74. DOI: 10.1126/science.1093857.

Vergin KL, Beszteri B, Monier A, Thrash JC, Temperton B, Treusch AH, Kilpert F, Worden AZ, Giovannoni SJ. 2013. High-resolution SAR11 ecotype dynamics at the Bermuda Atlantic Time-series Study site by phylogenetic placement of pyrosequences. The ISME journal 7:1322–32. DOI: 10.1038/ismej.2013.32.

Wang C, Malanotte-Rizzoli P. 2014. Diagnosis of physical and biological controls on phytoplankton distribution in the Sargasso Sea. Journal of Ocean University of China 13:32–44. DOI: 10.1007/s11802-014-1962-5.

Warwick-Dugdale J, Buchholz HH, Allen MJ, Temperton B. 2019a. Host-hijacking and planktonic piracy: How phages command the microbial high seas. Virology Journal 16:1–13. DOI: 10.1186/s12985-019-1120-1.

Warwick-Dugdale J, Solonenko N, Moore K, Chittick L, Gregory AC, Allen MJ, Sullivan MB, Temperton B. 2019b. Long-read viral metagenomics enables capture of abundant and microdiverse viral populations and their niche-defining genomic islands. PeerJ 7:e6800. DOI: 10.7717/peerj.6800.

Waterbury JB, Valois FW. 1993. Resistance to co-occurring phages enables marine Synechococcus communities to coexist with cyanophages abundant in seawater. Applied and Environmental Microbiology 59:3393–3399. DOI: 10.1128/aem.59.10.3393-3399.1993.

Wick RR, Holt KE. 2019. Benchmarking of long-read assemblers for prokaryote whole genome sequencing. F1000Research 8:2138. DOI: 10.12688/f1000research.21782.1.

Wilhelm W, Suttle C. 1999. Viruses and Nutrient Cycles in the Sea: Viruses play critical roles in the structure and function of aquatic food webs. BioScience 49:781–788.

Wilhelm LJ, Tripp HJ, Givan SA, Smith DP, Giovannoni SJ. 2007. Natural variation in SAR11 marine bacterioplankton genomes inferred from metagenomic data. Biology Direct 2:1–19. DOI: 10.1186/1745-6150-2-27.

Wilson WH, Mann NH. 1991. Lysogenic and lytic viral production in marine microbial communities.

Wright TD, Vergin KL, Boyd PW, Giovannoni SJ. 1997. A novel δ-subdivision proteobacterial lineage from the lower ocean surface layer. Applied and Environmental Microbiology 63:1441–1448. DOI: 10.1128/aem.63.4.1441-1448.1997.

Wu J, Sunda W, Boyle EA, Karl DM. Phosphate Depletion in the Western North Atlantic Ocean.

Zablocki O, Michelsen M, Burris M, Solonenko N, Warwick-Dugdale J, Ghosh R, Pett-Ridge J, Sullivan MB, Temperton B. 2021. VirION2: a short-and long-read sequencing and informatics workflow to study the genomic diversity of viruses in nature. PeerJ 9:e11088. DOI: 10.7717/peerj.11088.

Zhang Z, Qin F, Chen F, Chu X, Luo H, Zhang R, Du S, Tian Z, Zhao Y. 2020. Culturing novel and abundant pelagiphages in the ocean. Environmental Microbiology 00:1–17. DOI: 10.1111/1462-2920.15272.

Zhang L, Xu D, Huang Y, Zhu X, Rui M, Wan T, Zheng X, Shen Y, Chen X, Ma K, Gong Y. 2017. Structural and functional characterization of deep-sea thermophilic bacteriophage GVE2 HNH endonuclease. Scientific Reports 7:1–13. DOI: 10.1038/srep42542.

Zhao Y, Qin F, Zhang R. 2019. Pelagiphages in the Podoviridae family integrate into host genomes. Environmental microbiology 21:1989–2001. DOI: 10.1111/1462-2920.14487.

Zhao Y, Temperton B, Thrash JC, Schwalbach MS, Vergin KL, Landry ZC, Ellisman M, Deerinck T, Sullivan MB, Giovannoni SJ. 2013. Abundant SAR11 viruses in the ocean. Nature 494:357–60. DOI: 10.1038/nature11921.

Zhao Y, Wang K, Ackermann H-W, Halden RU, Jiao N, Chen F. 2010. Searching for a “hidden” prophage in a marine bacterium. Applied and environmental microbiology 76:589–595. DOI: 10.1128/AEM.01450-09.

