## Supplementary Information for "Long-read powered viral metagenomics in the Oligotrophic Sargasso Sea"

### Supplementary Methods

#### *Construction and identification of cellular metagenome assembled genomes*

To increase likelihood of host-prediction of phage contigs, the genomes of cellular Sargasso Sea microbes were assembled as follows: Short reads were mapped to the contigs from the cellular fraction using CoverM (v0.2.0) (https://github.com/wwood/CoverM). Reads were retained if they had > 95% identity and >75% read coverage with trimmed_mean used to remove the top 5% and bottom 5% depths. The bam files from read-mapping and the contigs from cellular samples were used as inputs into a custom script for binning. First, UniteM (v0.0.18) (UniteM; unpublished https://github.com/dparks1134/UniteM) was used to create a set of initial bins using the "unitem bin" command, utilizing the following binning options: gm2, bs, mb2, max40, max107, mb_verysensitive, mb_sensitive, mb_specific, mb_veryspecific. From this set of bins, UniteM "profile" and UniteM "consensus" commands were used to produce an ensemble bin set. Concordantly, DAS Tool (v1.1.1) (Sieber et al., 2018) and the MetaWRAP (v1.0.6) (Uritskiy, DiRuggiero & Taylor, 2018) "bin_refinement" module were utilized on the bins produced from the "unitem bin" command to produce ensemble bins. However, the MetaWRAP (v1.0.6) "bin_refinement" module only accepts three candidate bin sets, so MetaBAT2 (Kang et al., 2019), GroopM2 (Imelfort et al., 2014), and MaxBin2 (Wu, Simmons & Singer, 2015) outputs from UniteM were used as input into the MetaWRAP module. After all ensemble binning techniques were complete (MetaWRAP, DAS Tool, UniteM), the product ensemble bins were used as input for UniteM, MetaWRAP, and DAS Tool for a second iteration to produce an optimal bin set. Following the second iteration of ensemble binning, the output bins of each tool individually was evaluated for completeness and contamination using the CheckM (v1.0.12) (Parks et al., 2015) lineage workflow. Once the completeness and contamination statistics for the bin sets of the second iteration of ensemble binning tools were obtained, the bins greater than or equal to 70% completion and less than or equal to 10% contamination were used to calculate a quality score. Similar to the methods used in UniteM and CheckM, a score was calculated as the following: score = completeness – (2 x contamination). Each ensemble binning tool was scored, and then the tool with the highest quality score was used as the bin set for that particular sample. Following scoring, RefineM (v0.0.24) (Parks et al., 2017) “outliers” was used to remove any potential outliers associated with GC-content or tetranucleotide signatures with the following parameters: --gc_perc 95 –td_perc 95. The taxonomical classification of the resulting 89 MAGs was undertaken with GTDB-tk18 (Chaumeil et al., 2020); genomes were classified via placement in a GTDB reference tree.

#### ***Attempted host prediction***

Putative hosts were assigned to viral populations through prophage blast, tRNAscan-SE (v1.23) (Chen et al., 2019), and WisH (v1.0) (Galiez et al., 2017) using a scoring approach similar to what has been previously reported for human gut viromes (Gregory et al., 2020). In prophage blast, a nucleotide blast database was built by using Sargasso Sea MAGs. Viral representative contigs were used as input to BLAST against this database. Scores ranging from 1 to 4 were assigned based on percent identity and coverage (4: 98% ID and 90% cov, 3: 90% ID and 75% cov, 2: 90% ID and 50% cov, 1: 90% ID and 30% cov). General tRNA models were predicted for viral contigs in tRNAscan-SE (v1.23). Secondary structures of MAGs were searched using bacterial tRNA model. Scores were assigned to hits according to percent identity (3: 100%, 2: 95%, 1: 90%). Host models were built in WisH (v1.0). Null models were predicted by using 283 decoy RefSeq viral sequences that infect non-marine isolates belonging to the genera *Staphylococcus*, *Streptococcus*, *Lactobacillus*, *Propionibacterium*, *Mannheimia*, and *Paenibacillus* since none of these genera should encompass ocean MAGs. The virus-host linkages were predicted by providing target viral contigs, host MAG models, and the matrix of null model parameters. Scores were given based on reported p-values (p-value ≤ $10^{-10}$: 2.5, p-value ≤ $10^{-5}$: 2). Collectively, virus-host linkages that had scores ≥ 3 were considered as putative hosts.

### Supplementary Tables and Figures

| Statistic / Sample code | BS1 | BS3 | BS5 | 80m_unassigned |
|---|---|---|---|---|
| Mean read length (bp) | 4,100.1 | 3,840.1 | 3,738.8 | 3,927.9 |
| Mean read quality | 12.3 | 12.6 | 12.6 | 12.0 |
| Median read length (bp) | 3,662.0 | 3,534.0 | 3,441.0 | 3,583.0 |
| Median read quality | 12.6 | 12.8 | 12.8 | 12.3 |
| Number of reads | 363,245 | 481,694 | 509,715 | 2,399,558 |
| Read length N50 (bp) | 5,042.0 | 4,370.0 | 4,285.0 | 4,588.0 |
| Total bases (Mbp) | 1489.3 | 1849.7 | 1905.7 | 9425.2 |
| % reads >Q10 | 88.6 | 89.7 | 90.1 | 82.8 |
| Longest read >Q10 (bp) | 27114 | 20909 | 24884 | 32222 |

**Table S1.** Summary statistics for MinION long-read sequencing of Sargasso Sea viral communities (calculated with NanoStat; De Coster et al., 2018). Barcoded libraries for three samples taken from 80 m depth (encoded: BS1; BS3; BS5) were sequenced. Barcode ligation failed in >42% of reads; the sequencing depth remaining per sample was too low for successful downstream processing, so for further analysis sequences were pooled.

**Table S2.** Accession numbers (total: 438) of published cyanophage, pelagiphage and T4 genomes which were clustered to produce viral population representatives before recruitment of short reads recovered from Sargasso Sea viral- and cellular fraction samples.

See excel file: “cyanophage_pelagiphage_accession_numbers.txt”

**Table S3.** Metagenome Assembled Genomes (MAGs) generated from Sargasso Sea samples, including MAG Taxonomy, completion, and contamination.

See file: “BATS-MAGs-gtdbtk.summary”

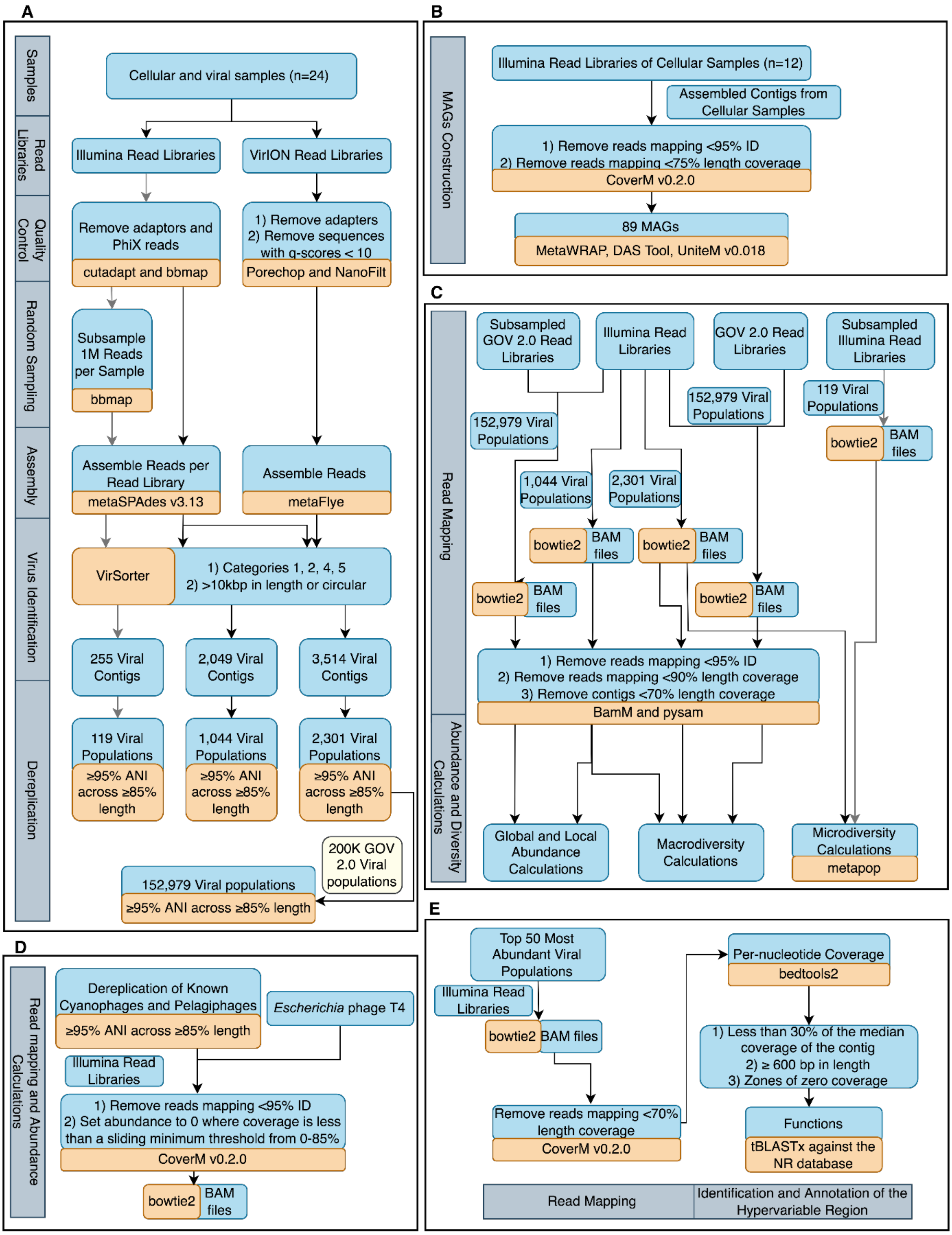


**Figure S1.** Schematic representation of the bioinformatic workflow for (A) the assembly and identification of viral populations, (B) the construction of MAGs, (C) the abundance and diversity calculation of viral populations, (D) the abundance calculation of known cyanophages and pelagiphages, and (E) viral Hypervariable Region (HVR) recovery and identification.

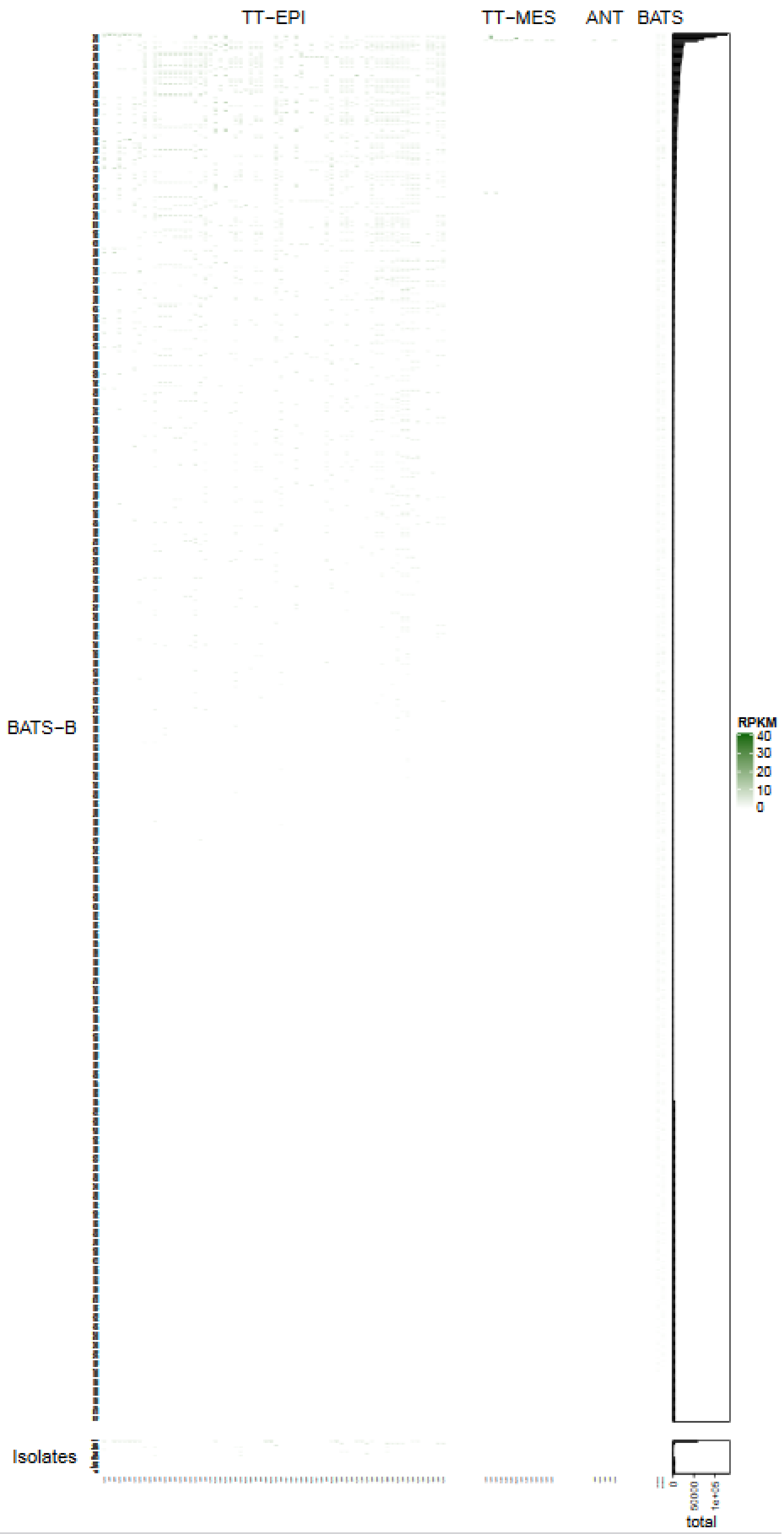
TT−EPI
TT−MES
ANT
BATS
BATS−B
Isolates
RPKM
40
30
20
10
0
total

**Figure S2.** SIMPER analysis showed that 674 Sargasso Sea viruses ('BATS-B', all of which were captured using long-read data) were important in discriminating the Sargasso Sea viromes from other viral communities/zones. Four members from these relatively rare viruses (bootstrapped median (n=10,000) number of Global Ocean Virome (GOV2) samples in which observed: 10 (4.5- 17 95% CI)) recruited a large number of reads from ≥1 temperate-tropical mesopelagic (TT-MES) or Antarctic (ANT) site implying some degree of viral import.

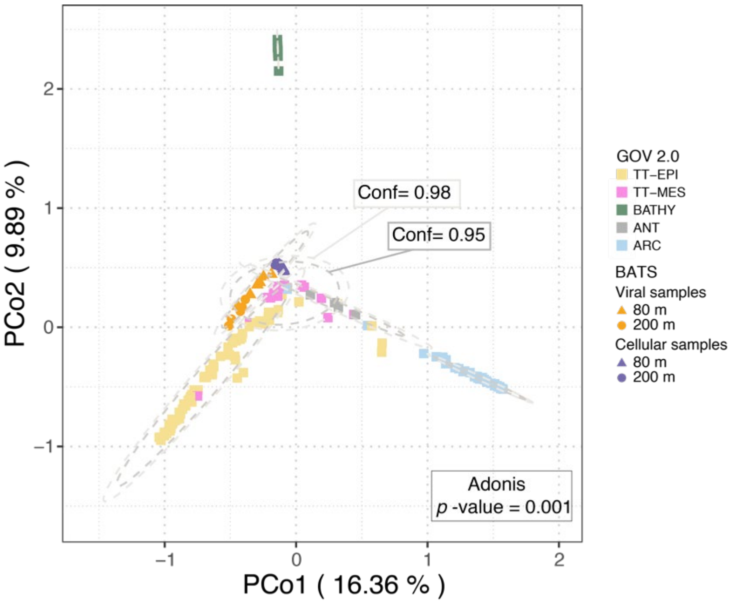


**Figure S3**. Principle coordinates analysis (PcoA) of a Bray-Curtis dissimilarity matrix calculated from mapping GOV 2.0 reads and BATS short reads to a combined dataset of BATS and GOV 2.0 viral populations with lengths greater than 10kbp. Viral community structure was suggested by ellipses drawn at 95% (inner) and 98% (outer) confidence intervals and analysis of variance (Adonis, p-value = 0.001). Three outlier GOV 2.0 viromes (station 155_SUR, station 72_MES, station102_MES; Gregory *et al.*, 2019) were not removed.

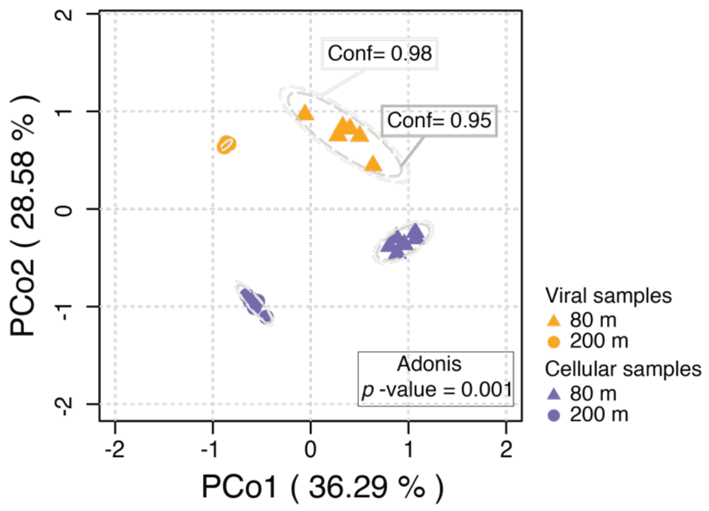


**Figure S4** Principal coordinates analysis (PcoA) of a Bray-Curtis dissimilarity matrix calculated from mapping BATS short reads to BATS viral populations (n = 1,044) derived from short-read assemblies. Viral community structure was suggested by ellipses drawn at 95% (inner) and 98% (outer) confidence intervals and analysis of variance (Adnois, p-value = 0.001).

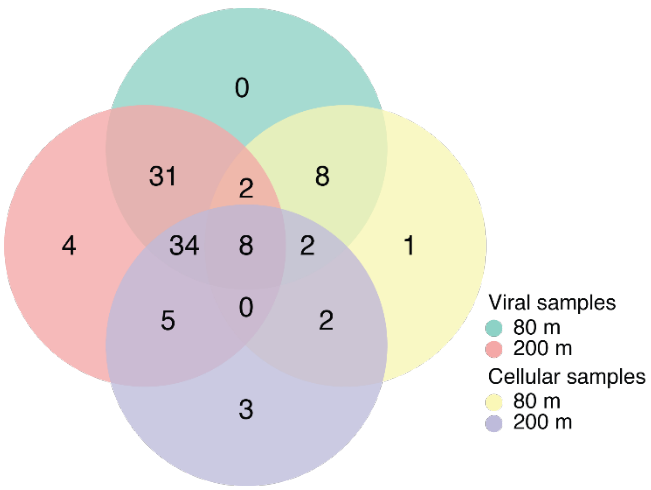


**Figure S5.** Top 100 most abundant Sargasso Sea viral populations identified from 1044 viral populations derived from short-read only assemblies: presence-absence in 80m viral samples, 80m cellular samples, 200m viral samples, and 200m cellular samples.

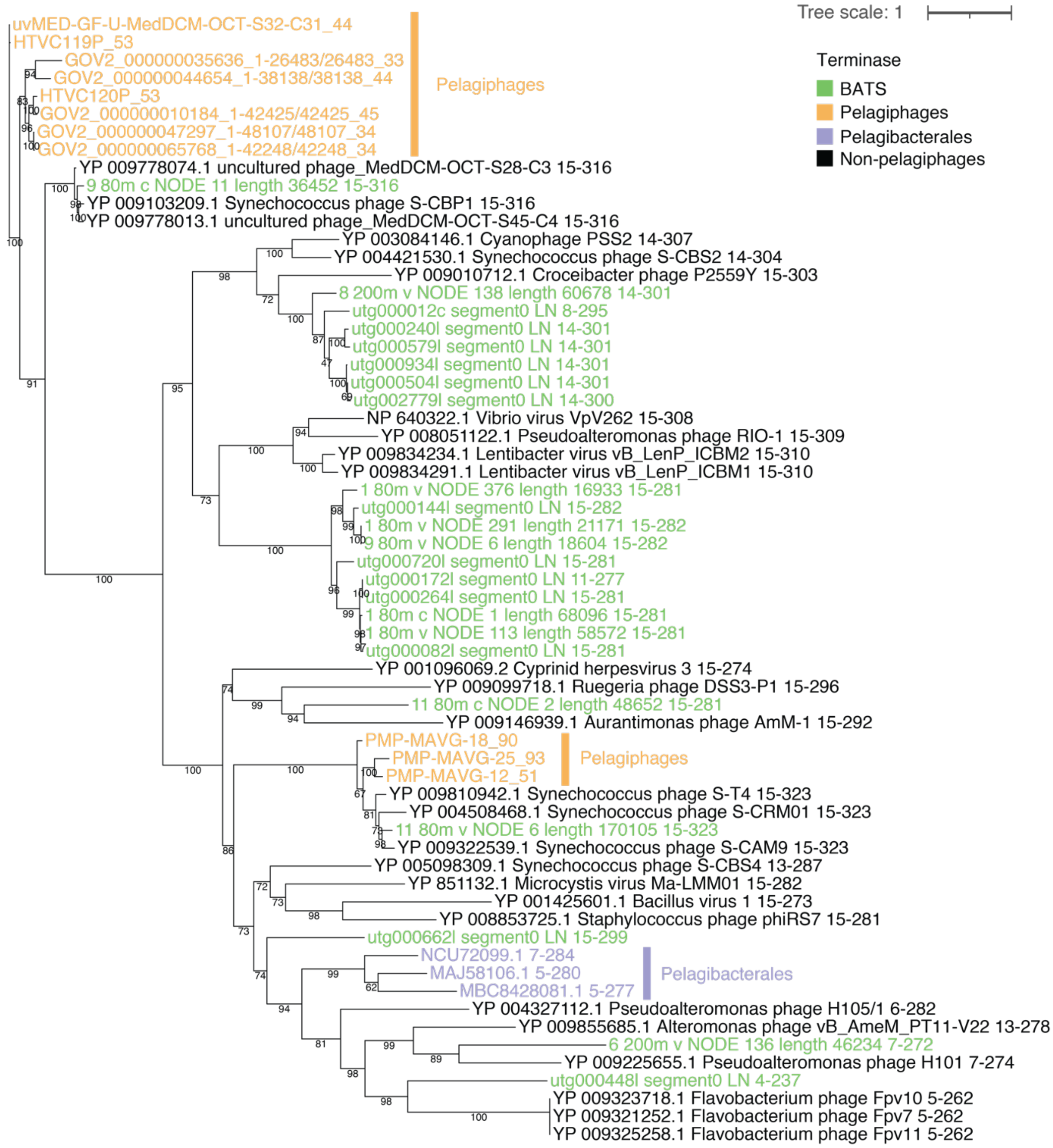


**Figure S6** Phylogenetic tree constructed using terminase (*TerL*) genes from published pelagiphage genomes, non pelagiphages, *Pelagibacterales* (identified via BLAST hits against terminase from known pelagiphages) and Sargasso Sea viral populations (*TerL* genes aligned using E-INS-i strategy; bootstrapping: 1000 iterations; clades subsampled for clarity).

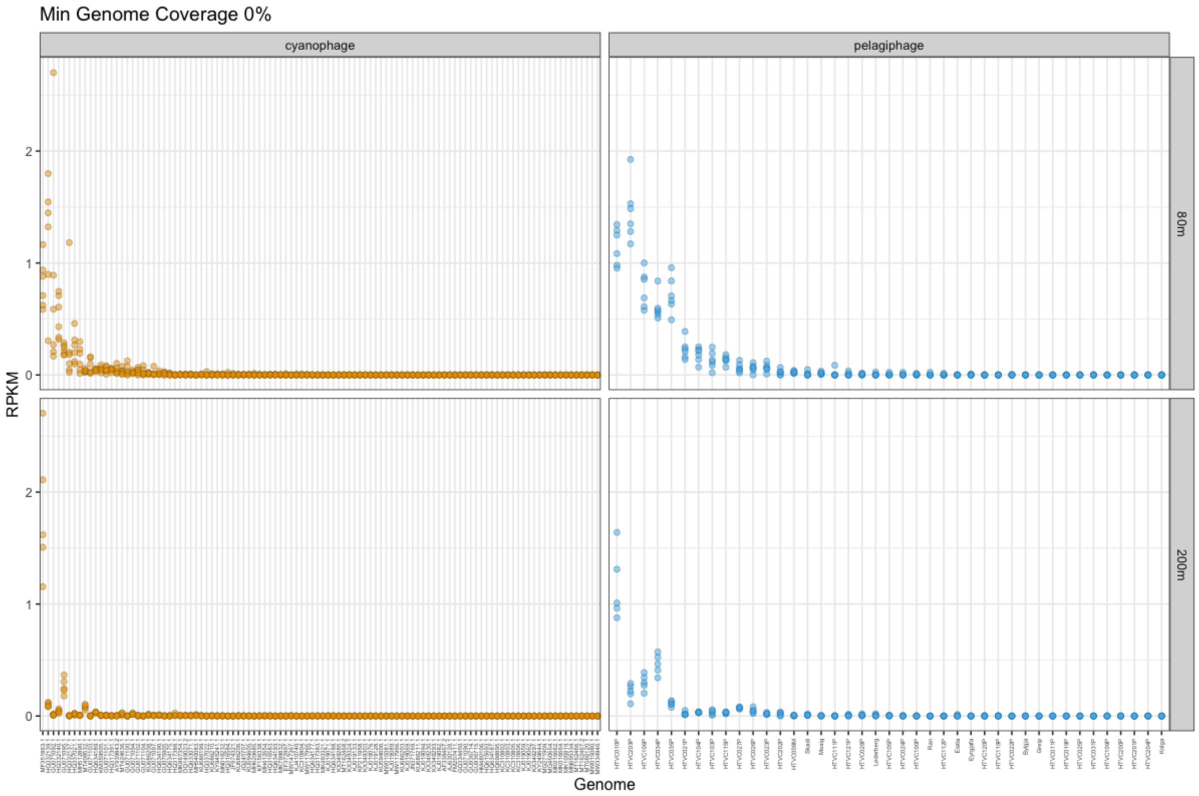

Min Genome Coverage 0%
cyanophage
pelagiphage
80m
200m
RPKM
Genome
0
1
2


**A**

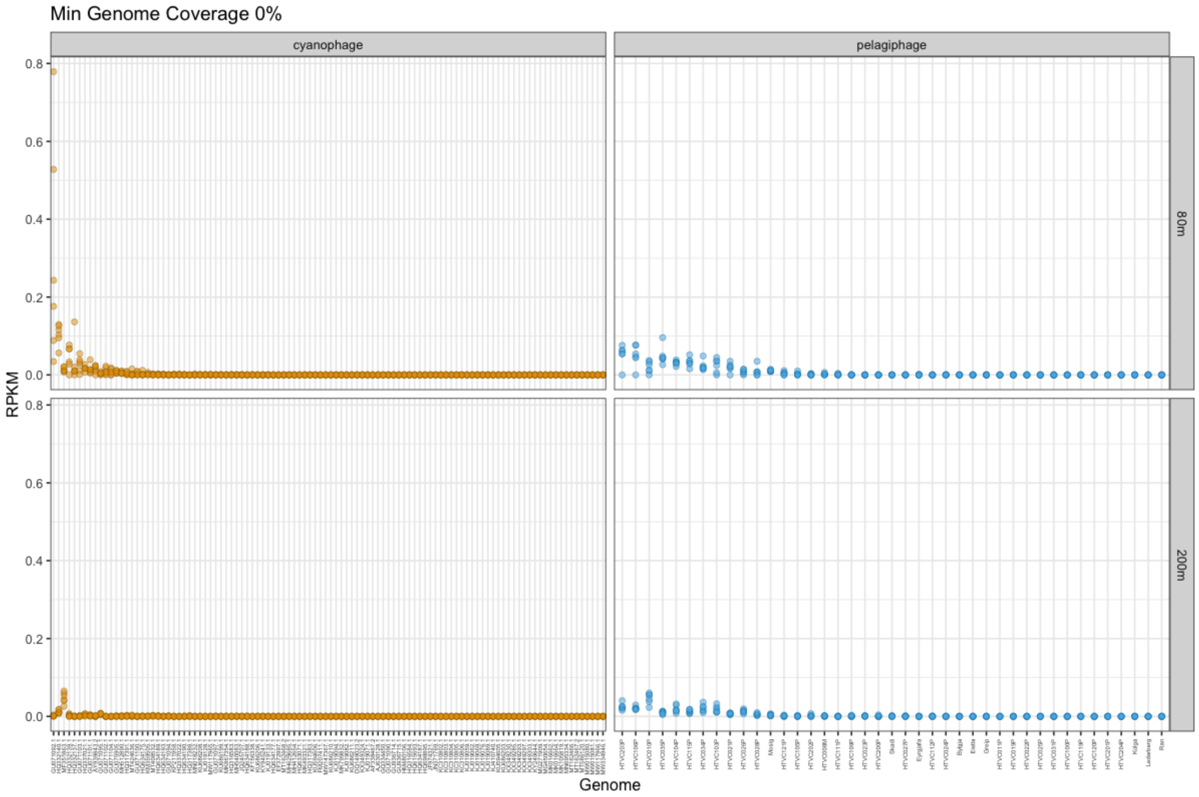

Min Genome Coverage 0%
cyanophage
pelagiphage
80m
200m
RPKM
Genome
0.0
0.2
0.4
0.6
0.8


**B**

**Figure S7.** Abundance (RPKM) of known cyanophages and pelagiphages in short-read data at 0-80% minimum genome coverage cut-off values in: **A.** the viral fraction; **B.** the cell-associated fraction. Cleaned Illumina sequences were competitively recruited (at ≥90% read length at ≥95% identity) to a dereplicated (at ≥ 95% nucleotide identity across ≥ 85% genome length) database of all published cyanophage and pelagiphage isolate genome (plus control *Escherichia* phage T4).

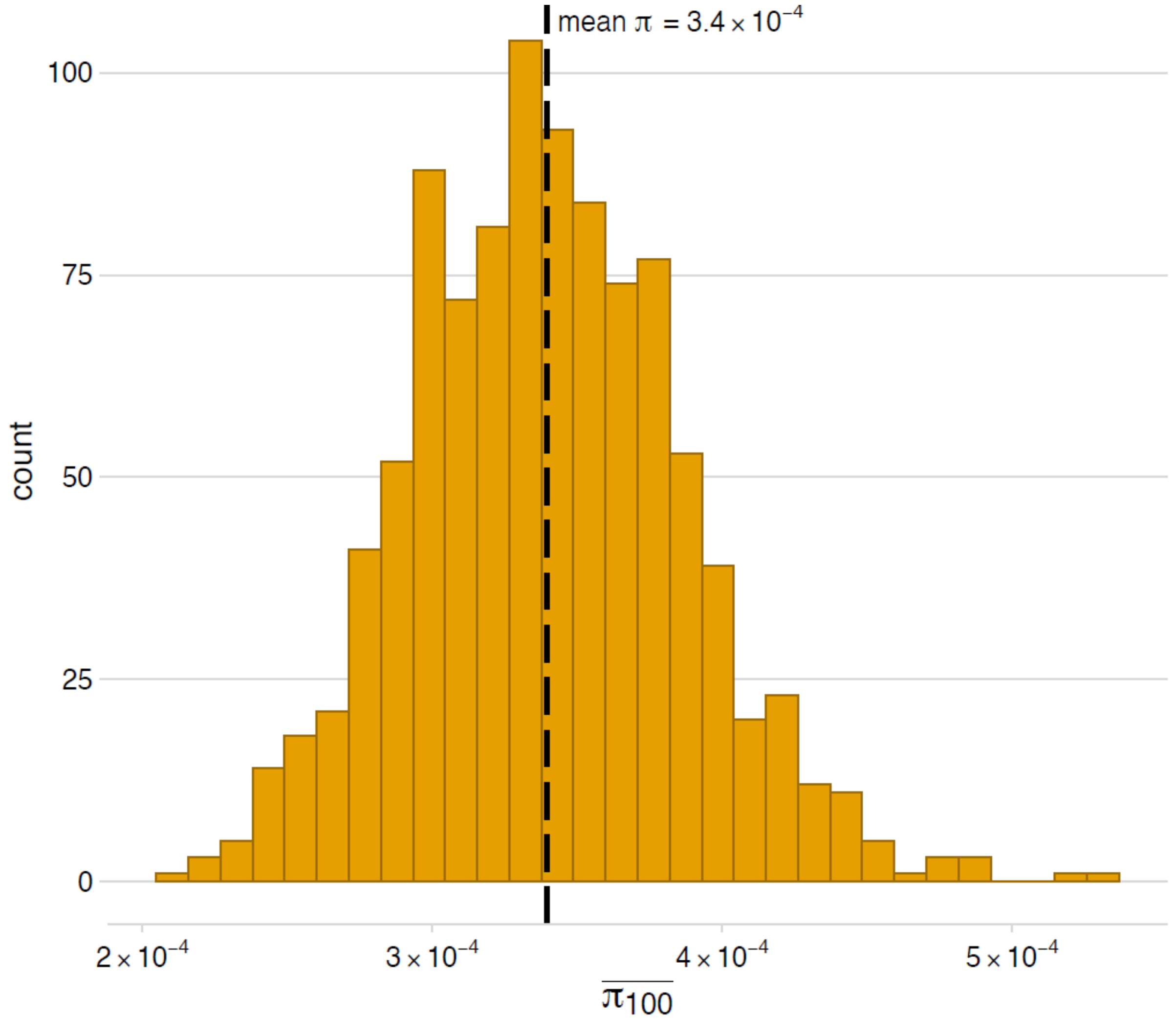


**Figure S8.** Average microdiversity (π) across Sargasso Sea viruses from both depths sampled (80m and 200m). Mean π was calculated as (Gregory et al., 2019): 100 π values were randomly subsampled from short--read Sargasso Sea viromes with replacement; the distribution mean π was generated via bootstrapping (1000 iterations), and 95% confidence intervals were calculated using the quantiles of this distribution.

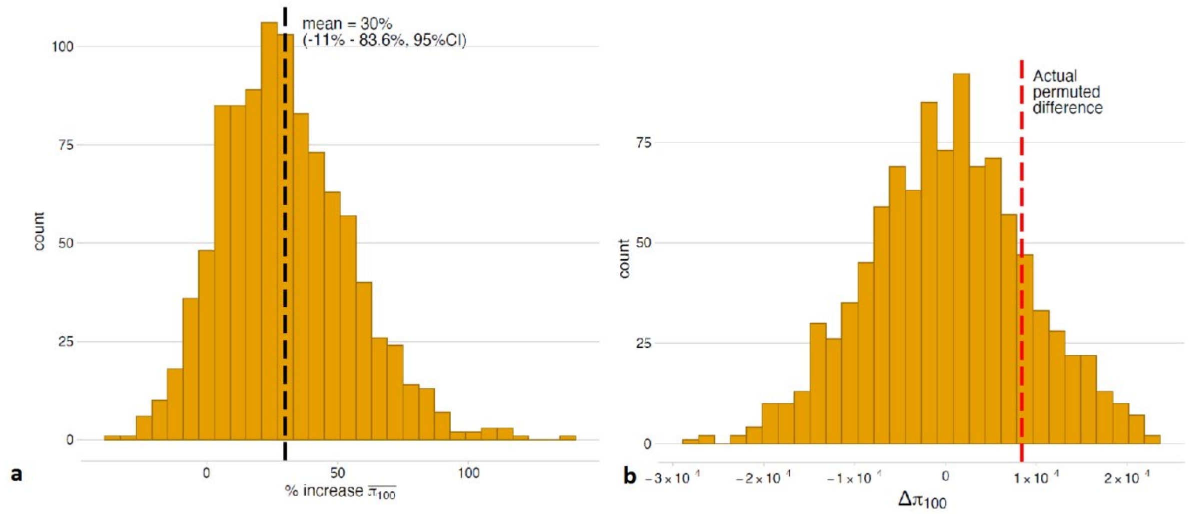


**Figure S9.** The permuted percentage increase of ~30% between average microdiversity values (mean π; calculated as (Gregory et al., 2019) of Sargasso Sea viruses from 80m and 200m (respectively) (**a**: 30.041% (-11.381 - 83.604, 95% CI), was not found to be significant (**b:** permuted significance test: $p$=0.164; under the null model, we observe a difference in the two populations at least as extreme as the actual measured difference (red line) 164 times out of 1000 permutations).

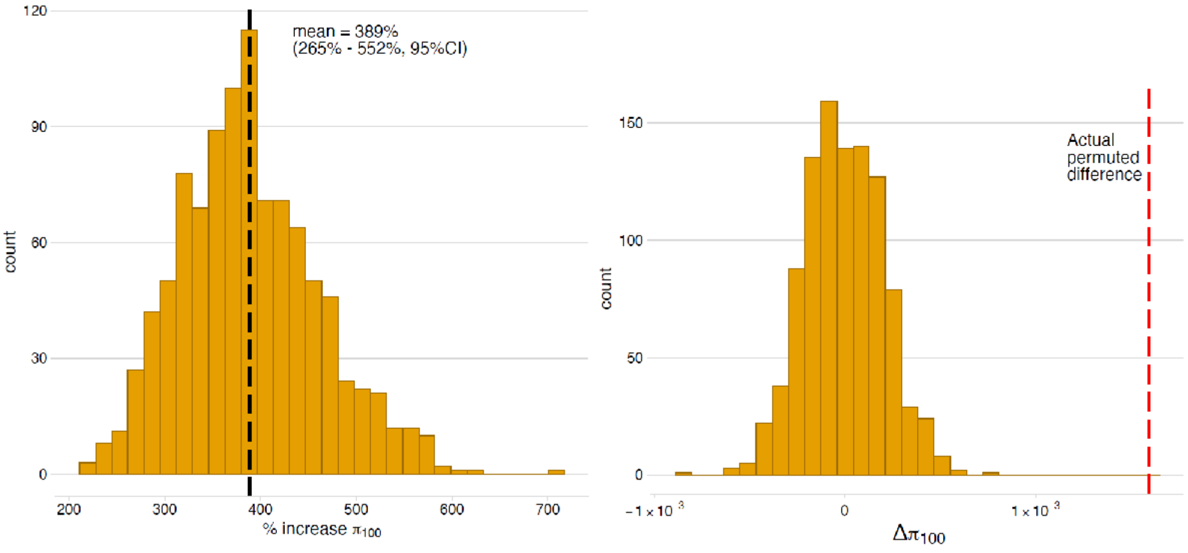


**Figure S10.** The permuted percentage increase of ~390% between average microdiversity values (mean π; calculated as (Gregory et al., 2019) Sargasso Sea viruses from 80m and 200m (respectively) (**a**: 388.668% (264.559 - 551.95%, 95% CI), was highly significant (**b:** permuted significance test: $p$ <0.001; under the null model, we never observe a difference in the two populations at least as extreme as the actual measured difference (red line) (0 times out of 1000 permutations).
