## Supplementary figures and images for "Long-read powered viral metagenomics in the Oligotrophic Sargasso Sea"

### Supplementary Figure S2

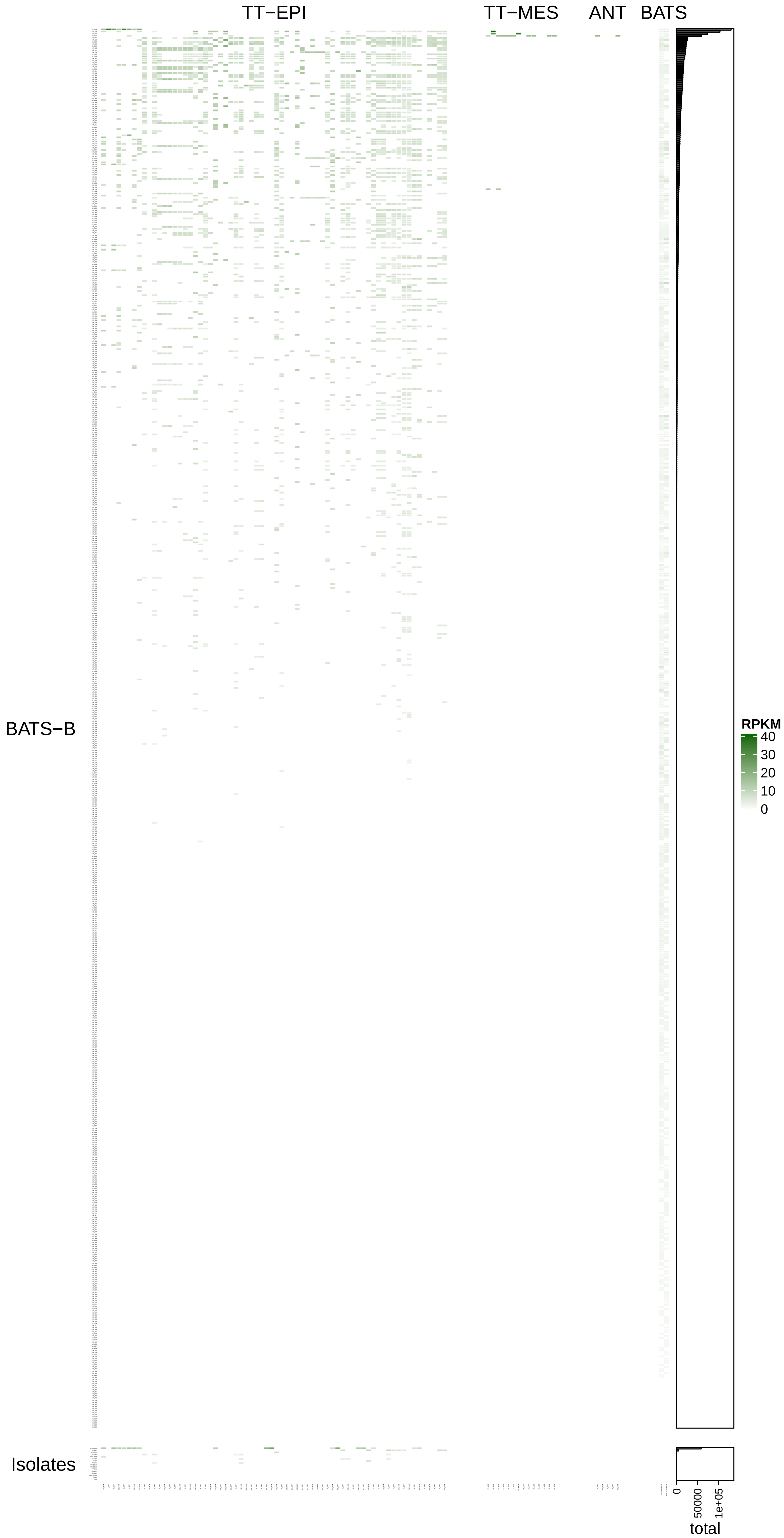

### Supplementary figure S7A

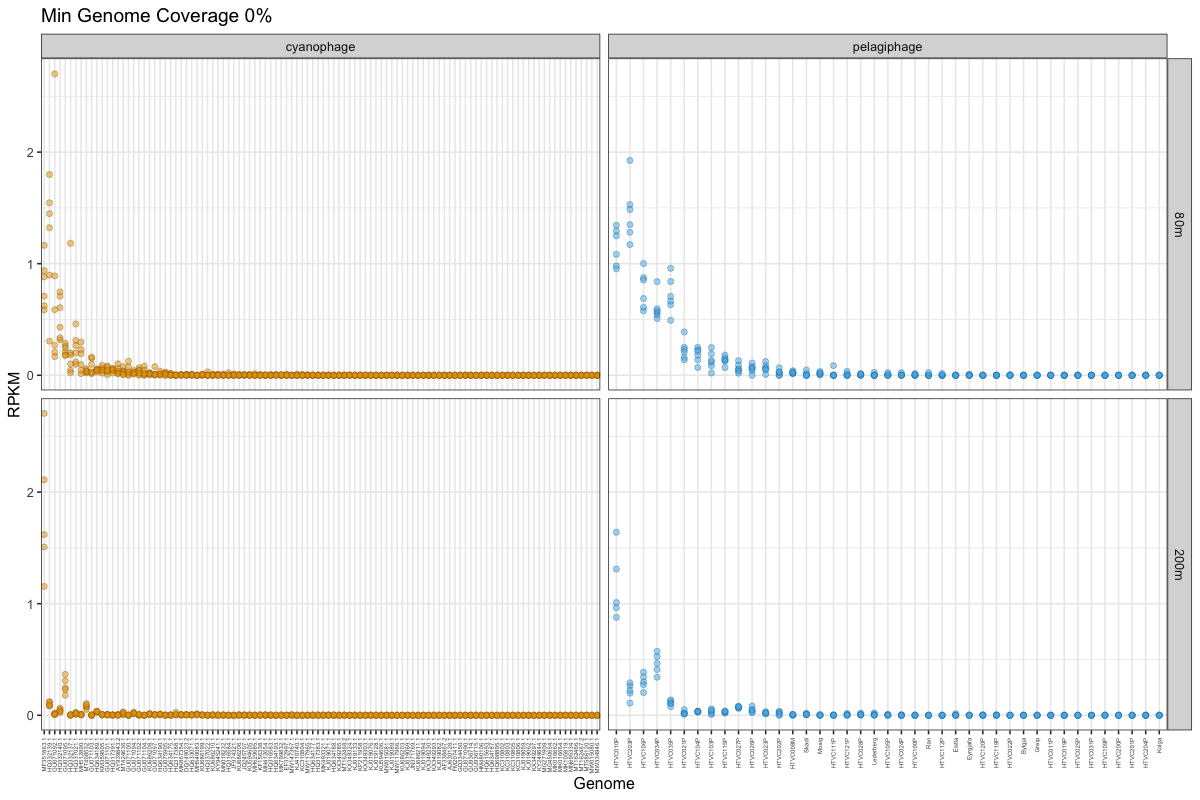

### Supplementary figure S7B

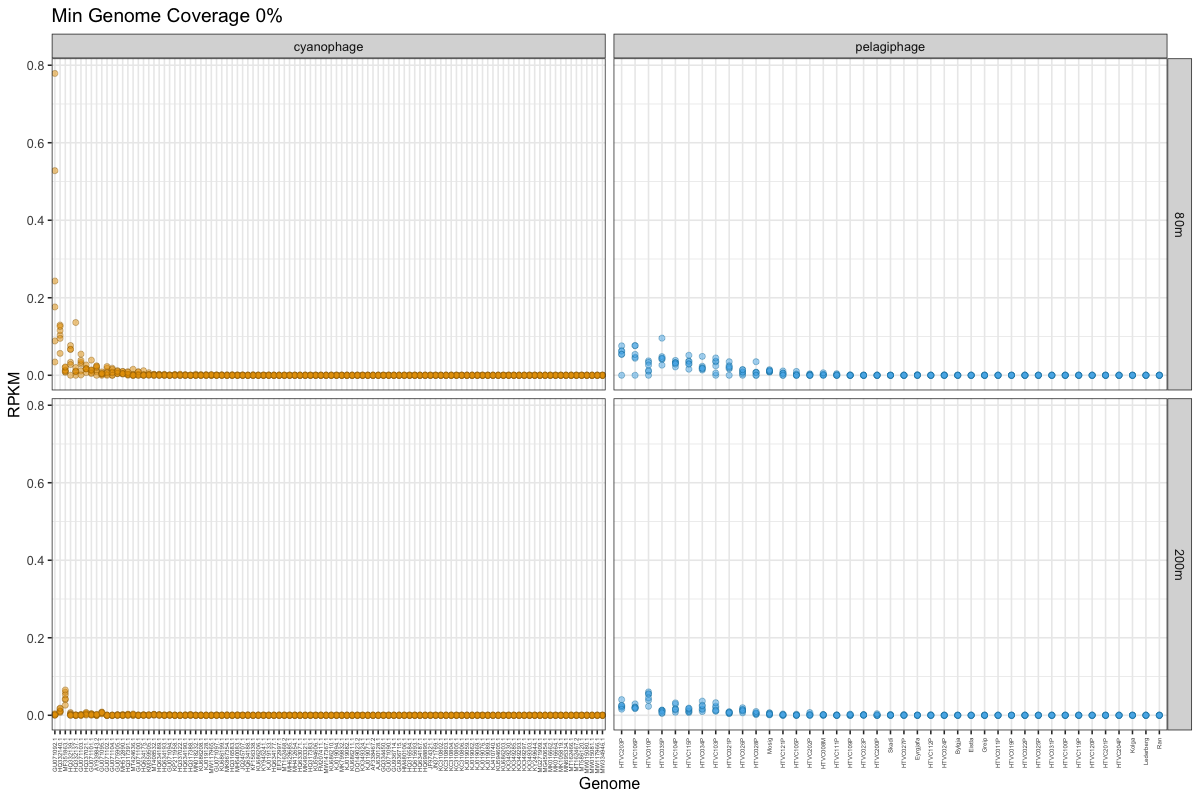
